# Geminin is required for *Hox* gene regulation to pattern the developing limb

**DOI:** 10.1101/2020.01.07.896472

**Authors:** Emily M.A. Lewis, Savita Sankar, Caili Tong, Ethan Patterson, Laura E. Waller, Paul Gontarz, Bo Zhang, David M. Ornitz, Kristen L. Kroll

## Abstract

Development of the complex structure of the vertebrate limb requires carefully orchestrated interactions between multiple regulatory pathways and proteins. Among these, precise regulation of 5’ *Hox* transcription factor expression is essential for proper limb bud patterning and elaboration of distinct limb skeletal elements. Here, we identified *Geminin* (*Gmnn)* as a novel regulator of this process. A conditional model of *Gmnn* deficiency resulted in loss or severe reduction of forelimb skeletal elements, while both the forelimb autopod and hindlimb were unaffected. 5’ *Hox* gene expression expanded into more proximal and anterior regions of the embryonic forelimb buds in this *Gmnn*-deficient model. A second conditional model of *Gmnn* deficiency instead caused a similar but less severe reduction of hindlimb skeletal elements and hindlimb polydactyly, while not affecting the forelimb. An ectopic posterior SHH signaling center was evident in the anterior hindlimb bud of *Gmnn*-deficient embryos in this model. This center ectopically expressed *Hoxd13*, the HOXD13 target *Shh*, and the SHH target *Ptch1*, while these mutant hindlimb buds also had reduced levels of the cleaved, repressor form of GLI3, a SHH pathway antagonist. Together, this work delineates a new role for *Gmnn* in modulating *Hox* expression to pattern the vertebrate limb.

**Summary:** This work identifies a new role for Geminin in mouse limb development. Geminin is a nuclear protein that regulates gene expression to control several other aspects of vertebrate development.

## Introduction

The vertebrate limb is a complex structure that develops during embryogenesis from a small bud into an appendage that is patterned along its anterior-posterior (A/P), dorsal-ventral (D/V), and proximal-distal (P/D) axes. Within each limb, three distinct skeletal compartments are formed, the stylopod or upper limb, zeugopod or lower limb, and the autopod or hand/foot, with each structure exhibiting asymmetry between the anterior (thumb) and posterior (little finger) sides. Limb development is a classically studied process for understanding embryonic patterning and tissue morphogenesis (reviewed in (Delgado and Torres, 2017; Petit et al., 2017; Sheeba et al., 2016; Tickle and Towers, 2017; Zuniga, 2015)). As experimental work in this area has advanced, it is evident that complex interactions between multiple regulatory molecules play key roles in patterning the limb. However, additional regulators of this process, and mechanisms by which it occurs, remain to be fully elucidated.

*Hox* genes encode transcription factors that play a central role in patterning and differentiation of multiple embryonic tissues and structures, both in the primary body axis and in the limb. The vertebrate genome contains four clusters of paralogous *Hox* genes (*HoxA-D*). Expression of genes in each *Hox* cluster is under complex control, involving both local and long-range chromatin structural changes (Andrey and Duboule, 2014; Montavon and Duboule, 2013; Montavon and Soshnikova, 2014; Neijts and Deschamps, 2017; Noordermeer and Duboule, 2013). *Hox* cluster gene expression exhibits spatial and temporal colinearity: 3’ located *Hox* genes in each cluster are expressed earlier and in more anterior embryonic locations, while 5’ genes are expressed later and more posteriorly (Andrey and Duboule, 2014; Izpisua-Belmonte et al., 1991; Kmita et al., 2002; Tarchini and Duboule, 2006; Tschopp and Duboule, 2011; Tschopp et al., 2009). 5’ located *HoxA* and *HoxD* cluster genes (*Hoxa9-a13* and *Hoxd9-d13*) are essential regulators of vertebrate limb development: *Hox9/10* paralogs have roles in stylopod, *Hox11* in zeugopod, and *Hox13* in autopod formation (Boulet, 2003; Davis et al., 1995; Fromental-Ramain et al., 1996a; Wellik and Capecchi, 2003). 5’ *Hox* gene expression also controls limb bud anterior-posterior polarity, promoting posterior-restricted expression of Sonic Hedgehog (*Shh*) to create the zone of polarizing activity (ZPA), which controls posterior limb bud patterning (reviewed in (Lopez-Rios, 2016; Tickle and Towers, 2017)). In the posterior limb bud mesenchyme, *Shh* expression is activated by binding of HOXD13 and HAND2 transcription factors to a distal *Shh* enhancer (Galli et al., 2010; Hill, 2007; Lettice et al., 2003; Sagai et al., 2005). Restriction of *Hoxd13* and *Hand2* expression to the posterior limb bud is controlled by GLI3, which undergoes constitutive proteolytic processing to a repressor form, GLI3R (te Welscher et al., 2002). During this period, limb bud outgrowth also occurs, and is centrally controlled by interactions between limb bud mesenchyme and FGF signaling from the overlying apical ectodermal ridge (AER) (Lewandoski et al., 2000; Niswander et al., 1993).

Expression of genes in the *Hox* clusters is controlled by epigenetic regulation, including histone methylation mediated by Polycomb complexes (PcG) (reviewed in (Andrey and Duboule, 2014; Montavon and Duboule, 2013; Montavon and Soshnikova, 2014; Neijts and Deschamps, 2017; Noordermeer and Duboule, 2013)). One of these PcG complexes tri-methylates histone H3 lysine 27 (H3K27me3) to repress gene expression, while locations co-enriched with this repressive H3K27me3 and with the active H3K4me3 modification are maintained in a ‘bivalent’ or poised expression state (Cao et al., 2002; Czermin et al., 2002; Kuzmichev et al., 2002; Muller et al., 2002). As in the primary body axis, PcG-catalyzed H3K27me3 restrains *Hox* expression in the developing limb. From mouse embryonic day 10.5 (E10.5), H3K27me3 is cleared from 5’ *Hox* genes in the posterior limb bud mesenchyme to promote their expression (Montavon and Duboule, 2013; Soshnikova and Duboule, 2009; Williamson et al., 2012). Activation of 5’ *Hox* gene expression at this time also involves large-scale chromatin reorganization and long-range chromatin looping between the 5’ *Hox* gene cluster and multiple enhancers (Andrey and Duboule, 2014; Andrey et al., 2013; Gonzalez et al., 2007; Montavon et al., 2011; Spitz et al., 2003).

As a central regulator of developmental genes, PcG activity is likewise modulated by interactions with other regulatory proteins, including Geminin (GMNN), a nuclear protein that can interact directly with PcG complexes (Luo et al., 2004). In multiple developmental contexts, physical and/or functional interactions between GMNN and PcG modulate its regulation of developmental gene expression. GMNN cooperates with PcG to restrain *Hox* expression during primary body axis patterning and to control mesendodermal specification of pluripotent embryonic cells (Caronna et al., 2013; Lim et al., 2011; Luo et al., 2004). GMNN also represses *Hox* expression during later developmental events, such as hematopoietic stem cell differentiation (Karamitros et al., 2014). In addition to working cooperatively with PcG complex activity to restrain *Hox* expression in several developmental contexts, GMNN can also interact directly with a subset of the HOX transcription factors, and may negatively regulate their activity (Salsi et al., 2009; Zhou et al., 2012). Models of deficiency in the mouse have revealed roles for *Gmnn* in early embryogenesis, in specification and patterning of the forming neural tube and paraxial mesoderm, and in several aspects of stem cell differentiation and later tissue patterning (Barry et al., 2012; Emmett and O’Shea, 2012; Gonzalez et al., 2006b; Karamitros et al., 2015; Ma et al., 2016; Patterson et al., 2014). However, a role for *Gmnn* in limb development has not previously been identified. Here, we uncovered novel requirements for *Gmnn* in several aspects of embryonic limb bud patterning. We demonstrated that, in these models, 5’ *Hox* gene expression in the limb bud is altered, and appears to be a major contributor to the limb developmental defects observed.

## Results

### Loss and reduction of forelimb stylopod- and zeugopod-derived skeletal elements in the *Gmnn^f/f^*; Prx-Cre model of Geminin deficiency

To identify requirements of *Gmnn* for embryonic patterning, we crossed a *Gmnn* floxed allele with several Cre lines that drive excision both in developing mesoderm of the primary body axis and in the limb mesenchyme, with different temporal and spatial profiles. Prx1-Cre [Tg(Prrx1-cre)1Cjt] expresses Cre recombinase under the control of a *Prrx1* enhancer. The transgene is strongly expressed in forelimb bud mesenchyme from embryonic day 9.5 (E9.5), with only weak expression in single cells of the hindlimb ridge mesenchyme at this stage (Logan et al., 2002). At E10.5, the transgene continues to be strongly expressed in forelimb mesenchyme, while we still detected only weak expression in hindlimb mesenchyme at this stage (**Fig. S1**). At this time, flank, cranial, and craniofacial tissues of the embryo also express Prx1-Cre (Logan et al., 2002).

To avoid germline Cre recombinase activity observed in females but not males, *Gmnn^+/f^;* Prx-Cre males were mated to *Gmnn^f/f^* females. *Gmnn^f/f^*; Prx-Cre mutant animals were obtained from these crosses at Mendelian ratios (25.23%; n=107; **Table S1A**), and developed into adults. While development of the axial skeleton and hindlimbs appeared normal in these animals, they exhibited specific forelimb defects (**Fig. 1**). We scored these forelimb phenotypes in skeletal preparations from *Gmnn^f/f^*; Prx-Cre mutant animals (n=10) versus *Gmnn^+/f^*; Prx-Cre littermate controls (n=6) (**Table S1B-C**). None of the forelimbs in mutant animals were normal, with two-thirds (70%) of the animals examined missing the humerus, the stylopod-derived forelimb element, and either the radius, ulna, or both of these zeugopod-derived skeletal elements, either in one or both forelimbs. The most mildly affected animals (30%) retained all skeletal elements, but these were shortened and/or of reduced size. However, both the forelimb autopod and the hindlimbs appeared normal in these animals.

**Figure 1.**
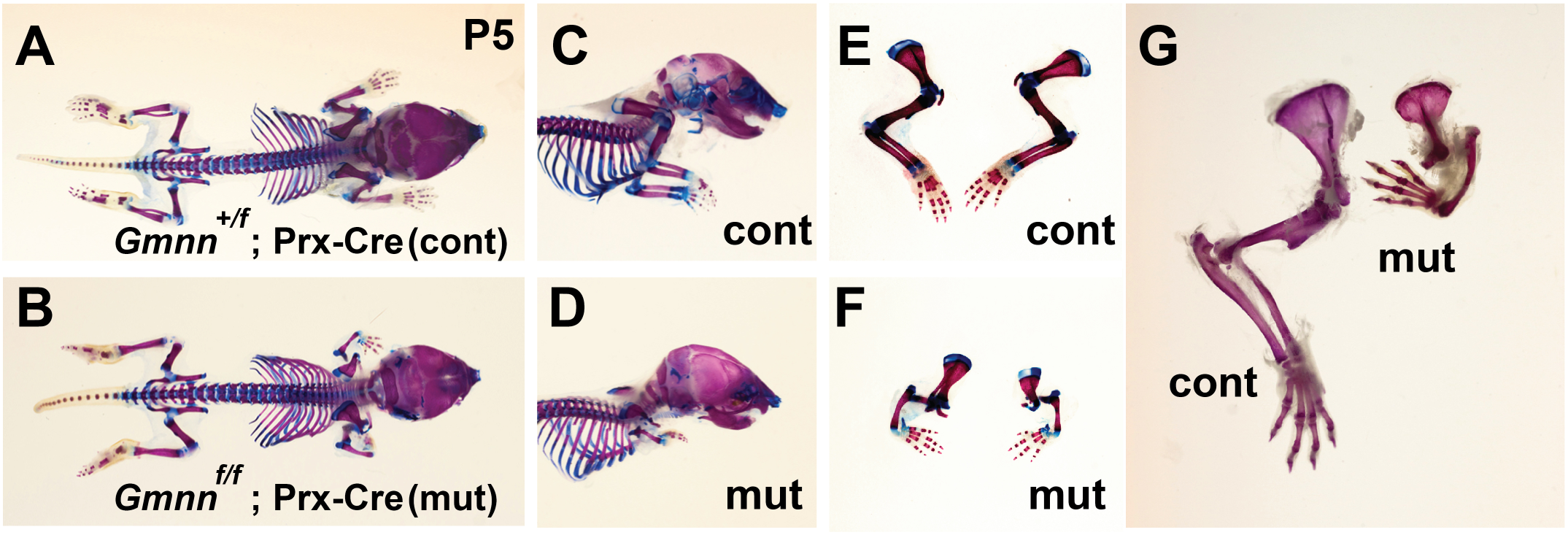
Loss or reduction of stylopod- and zeugopod-derived forelimb skeletal elements in the *Gmnn^f/f^*; Prx-Cre conditional model of Geminin deficiency. Skeletal preparations were performed on post-natal day 5 (P5) animals with the *Gmnn^+/f^*; Prx-Cre (control) and *Gmnn^f/f^*; Prx-Cre (mutant) genotypes. (A-B) Dorsal and (C-D) side views of P5 animals, and (E-G) isolated limb views of the forelimbs are shown. A summary of findings, including numbers of animals analyzed, is provided in Table S1.

We further analyzed when this skeletal phenotype was evident during embryogenesis by collecting embryos at E14.5, E16.5 and E18.5, and examining hematoxylin and eosin stained sections through the developing forelimbs of mutant and control embryos. By E14.5, some skeletal elements were already missing or reduced in size, while the structure of those that did form did not appear to be grossly disrupted, by comparison with controls (**Fig. S2**). As loss or reduction of skeletal elements was already apparent by E14.5, disruption of earlier specification and patterning processes in the forelimb bud appeared be a major driver of the phenotypes observed.

### Hindlimb polydactyly and skeletal element reduction in the *Gmnn^f/f^*; *Dermo^Cre^* model of Geminin deficiency

We compared these phenotypes with those that resulted from crossing this *Gmnn* conditional allele with a different Cre driver that is also expressed in the limb bud mesenchyme, *Dermo^Cre^*. *Dermo^Cre^* [Twist2^tm1(cre)Dor^] is a Cre recombinase knock-in, replacing exon 1 of the *Twist2* gene (Yu et al., 2003). Recombinase activity is detectable from E9.5 at the surface of the embryo and in mesenchymal condensations that contribute to both the axial and limb skeletal elements (**Fig. S1**) (Yu et al., 2003).

*Gmnn^f/f^; Dermo^Cre^* animals were also born in Mendelian ratios (24.6%; n=130), and grew into fertile adults (**Table S2A**). While the primary body axis and forelimbs of all mutant embryos appeared phenotypically normal, the animals exhibited specific hindlimb defects. Polydactyly was apparent in the hind feet (92.3%; n=26; **Table S2B-C**), with most animals having 6-7 toes and with syndactyly visible in some at adult stages (**Fig. 2**). When bone preparations were generated from animals at E18.5 up to post-natal day 5, the other hindlimb skeletal elements appeared grossly normal. However, skeletal preparations from adults revealed shortening and curvature of both the hindlimb stylopod-(femur) and zeugopod-(tibia/fibula) derived skeletal elements in most animals (87.5%; n=16; **Fig. 2**; **Table S2B-C**). This was reminiscent of the shortening and curvature seen in the forelimbs of Prx-Cre mutant animals, although substantially less severe, while complete loss of any hindlimb skeletal elements (the tibia/fibula) was only apparent in one animal (3.8%; n=26).

**Figure 2.**
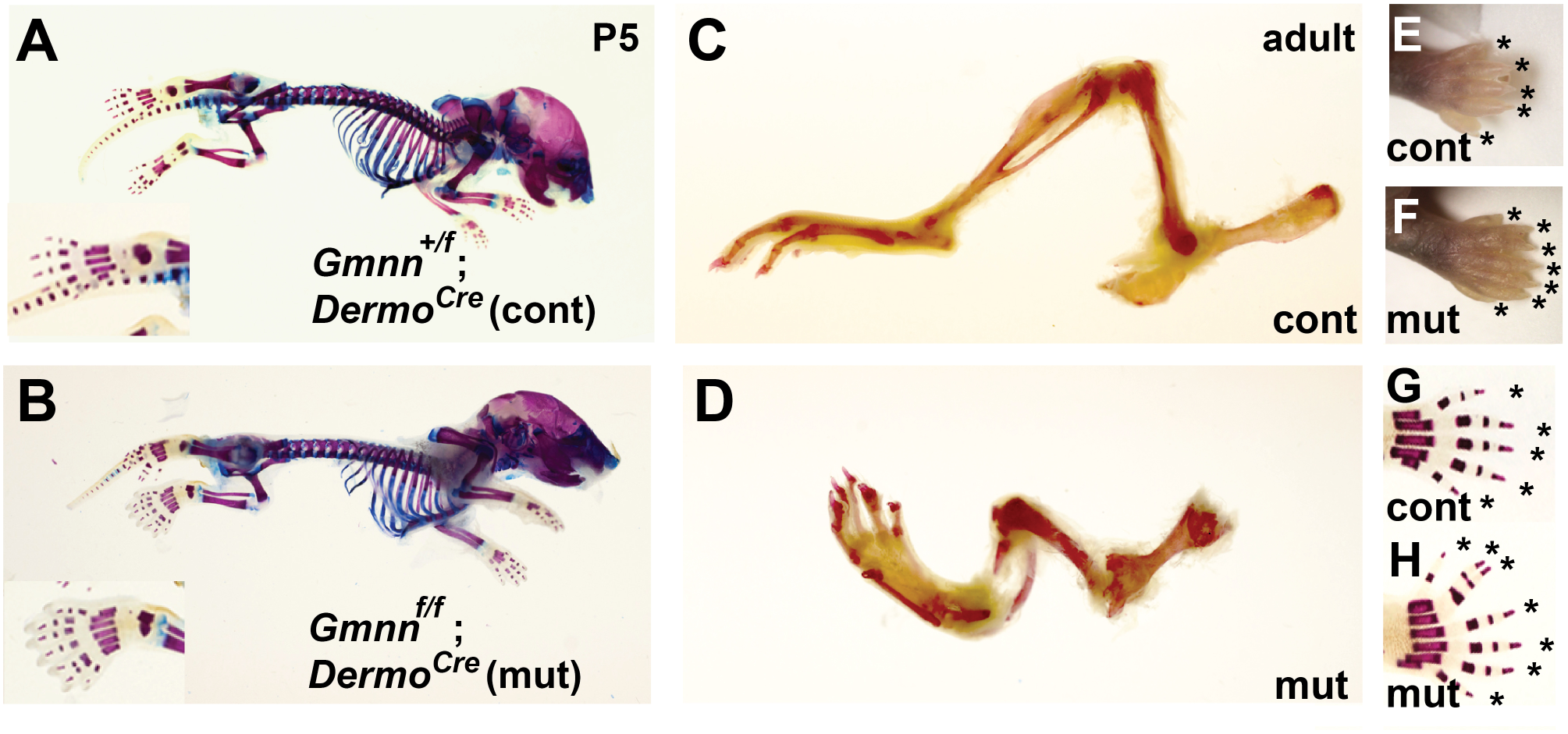
Hindlimb polydactyly and reduction and abnormality of skeletal elements in the *Gmnn^f/f^; Dermo^Cre^* conditional model of Geminin deficiency. Skeletal preparations were performed on animals with the *Gmnn^+/f^*; *Dermo^Cre^* (control) or *Gmnn^f/f^; Dermo^Cre^* (mutant) genotypes at P5 (A-B, E-H) and adult (C-D) stages. (A-B) Side view of animal, with hindlimb autopod shown in inset. (C-D) Isolated hindlimb from adult. (E-H) P5 hindlimb autopod (E-F) images and (G-H) bone preparations with * indicating autopod digits. A summary of findings, including numbers of animals analyzed, is provided in Table S2.

### Phenotypic consequences of Geminin deficiency are dependent on timing and location

Since *Gmnn^f/f^*; Prx-Cre and *Gmnn^f/f^; Dermo^Cre^* mutant animals respectively exhibited fore-versus hindlimb specific skeletal defects, we assessed whether this could be related to differences in the spatial and temporal expression of *Gmnn*. We initially examined *Gmnn* expression in the limb bud by using the *Gmnn* allele in its ‘knock-out first’ configuration, in which introduction of a FRT-site flanked *β-galactosidase* reporter into the *Gmnn* locus both disrupts the gene and reports on *Gmnn* expression. Based upon *β-galactosidase* reporter expression in E10.5-12.5 embryos and in limb buds heterozygous for this allele, *Gmnn* appears to be broadly expressed in both the embryo and in the developing limb buds, consistent with prior work documenting *Gmnn* expression in many proliferating cell populations during embryonic development (Emmett and O’Shea, 2012; Gonzalez et al., 2006a; Patterson et al., 2014) (**Fig. S3**). As we did not detect overt differences in *Gmnn* expression between the developing fore- and hind-limb buds, we next compared the spatial and temporal expression of the Prx-Cre and *Dermo^Cre^* drivers in the developing limb buds. We crossed both Prx-Cre and *Dermo^Cre^* strains to the R26R reporter line and examined *LacZ* expression in the limb buds from E9.5. As described previously, Prx-Cre was strongly expressed in the forelimb bud by E9.5, while no expression in cells of the hindlimb ridge could be detected at this time (Logan et al., 2002). By E10.5, Prx-Cre was strongly expressed in the forelimb bud, but only weakly expressed in the hindlimb bud (**Fig. S1**). By comparison, *Dermo^Cre^* expression could not be detected in the forelimb bud at E9.5, while it was expressed in some cells of the hindlimb ridge at this stage. By E10.5, *Dermo^Cre^* was strongly expressed in both the fore- and hindlimb bud (**Fig. S1**). Early aspects of limb bud patterning occur approximately a half day later in the hindlimb bud than the forelimb bud (Lopez-Rios, 2016). As *Dermo^Cre^* expression in the hindlimb bud from E9.5 and in both the fore- and hindlimb buds from E10.5 altered hindlimb but not forelimb development, while Prx-Cre excision from E9.5 in the forelimb bud instead only altered forelimb development, these differential effects suggest a window of sensitivity to *Gmnn* deficiency corresponding to early events of limb bud outgrowth and patterning.

Interestingly, although *Gmnn* was excised from E9.5 or E10.5 in some mesodermal tissues in the primary body axis by both Prx-Cre and *Dermo^Cre^* drivers (**Fig. S1**), the adult animals had normal axial skeletal patterning. Therefore, to determine whether *Gmnn* was required in primary body axis mesoderm at earlier developmental stages, we crossed the same floxed *Gmnn* allele to a pan-mesodermal driver, T-Cre, which drives Cre-mediated recombination in gastrula mesoderm from E7.5 (Perantoni et al., 2005). We were unable to recover embryos with the mutant genotype at stages later than E9.5 while, at this stage, *Gmnn^f/f^*; T-Cre embryos were smaller in size than littermate controls and had relatively normal head and rostral trunk tissue patterning, but either tissue deficiency or gross abnormality in the caudal trunk and tail regions (**Fig. S4A-B**). We assessed whether either altered development or differences in proliferation or apoptosis could contribute to this caudal phenotype by performing immunohistochemistry on sections of mutant embryos and littermate controls at E8.5, an earlier developmental timepoint prior to turning (**Fig. S4C-O**). Markers of the three germ layer derivatives and of proliferating cells were expressed similarly in mutant and control embryos (**Fig. S4E-M**). However, cleaved Caspase, a marker of apoptosis, exhibited increased expression in sections through the caudal end of the embryo (**Fig. S4N-O**). Therefore, increased apoptosis, particularly at the caudal end of the embryo, appears to contribute to the tissue deficiencies apparent in *Gmnn^f/f^*; T-Cre mutant embryos by E9.5 (**Fig. S4**). The axial patterning defects and embryonic lethality observed in the *Gmnn^f/f^*; T-Cre mutant embryos support a requirement for *Gmnn* in gastrula and post-gastrula mesoderm, while induction of *Gmnn* deficiency in some mesodermal tissues of the later embryo, by using either the *Dermo^Cre^* or Prx-Cre drivers, did not result in axial patterning defects.

### *Gmnn^f/f^*; Prx-Cre embryos exhibit altered expression of 5’ *HoxD* cluster genes

We next explored the molecular basis of the *Gmnn^f/f^*; Prx-Cre limb phenotype by collecting embryos during limb bud outgrowth and patterning and examining markers of limb anterior-posterior patterning, proximo-distal outgrowth, and regional specification. The finding that *Gmnn^f/f^*; Prx-Cre forelimbs exhibited loss or reduction of skeletal elements suggested that misregulation of the 5’ *HoxD* cluster genes (*Hoxd10-d13*) during limb bud patterning could contribute to the defects seen. 5’ *HoxA/D* cluster genes are required for specific aspects of limb regional patterning, with roles for *Hox9/10, Hox11*, and *Hox13* paralogs in the formation of stylopod-, zeugopod-, and autopod-derived structures, respectively. During limb bud outgrowth, the 5’ *Hox* genes are expressed in a nested pattern that parallels their requirement for proper development of different proximo-distal regions, with *Hox10-Hox13* paralogs exhibiting progressively less to more distally restricted expression, while *Hox10-12* paralogs likewise exhibit lower to higher anterior-posterior enrichment of expression in the limb bud (**Fig. 3**). We examined *Hox* gene expression by whole mount in situ hybridization (WISH), performing each in parallel WISH analysis on pairs of somite-matched embryos obtained from the same timed pregnancy with *Gmnn^f/f^*; Prx-Cre mutant versus *Gmnn^+/f^*; Prx-Cre control genotypes, respectively. In forelimb buds of *Gmnn^f/f^;* Prx-Cre mutant embryos, we found that the normal pattern of *Hox* expression was disrupted, with expression of multiple 5’ *Hox* genes including H*oxd10, Hoxd11, Hoxd12, Hoxd13*, and *Hoxa13* expanding both into more proximal and more anterior regions of the forelimb bud (**Fig. 3**; **Table S3A**).

**Figure 3.**
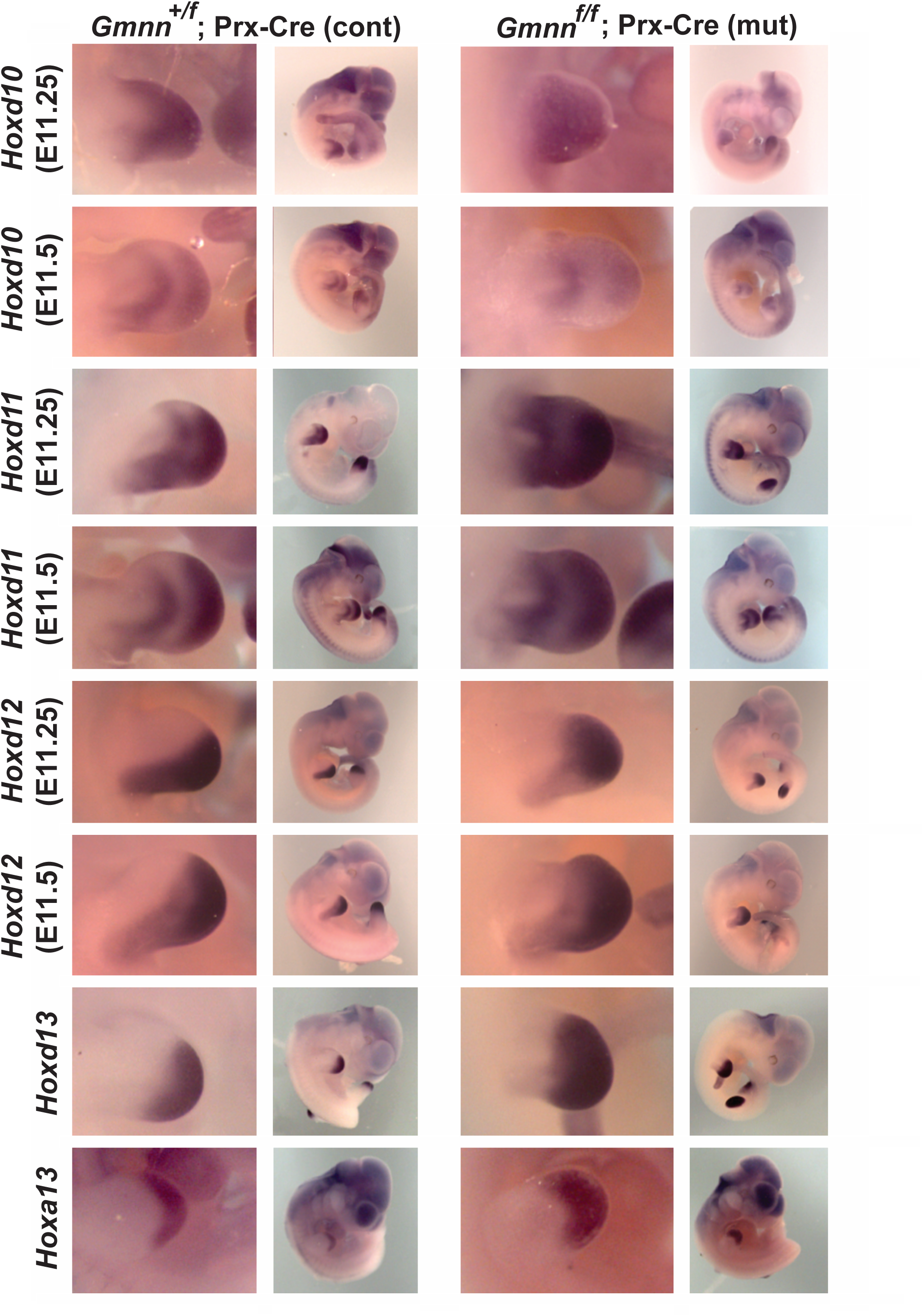
Forelimb buds of *Gmnn^f/f^*; Prx-Cre mutant embryos exhibit expanded domains of 5’ *Hox* gene expression. Pairs of *Gmnn^+/f^*; Prx-Cre (control) and *Gmnn^f/f^*; Prx-Cre (mutant) somite-matched embryos obtained from the same timed pregnant female were used for whole mount in situ hybridization with probes specific to the genes indicated. For Hoxd10-12, two examples are shown at slightly younger (E11.25) and older (E11.5) stages, to capture the dynamic changes in normal *Hox* expression patterns that occur at this time. A summary of findings, including number of biological replicates for each WISH probe, is provided in Table S3A.

We also examined several other classes of markers in *Gmnn^f/f^*; Prx-Cre mutant versus controls. Of the posterior limb bud markers assessed, *Ptch1* exhibited a slight anterior expansion of its expression domain, while *Hand2* and *Shh* expression were similar in mutant and control embryos and ectopic anterior *Shh* expression was not observed, suggesting that limb bud patterning along the anterior/posterior axis patterning was largely unaffected (**Fig. 4**). Expression of *Fgf8*, which marks the apical ectodermal ridge (AER) and is a central driver of proximo-distal limb outgrowth, was similarly expressed in mutant and control embryos (**Fig. 4**). We additionally assessed markers of proliferating (Ki67), mitotic (phosphorylated histone H3; pH3), and apoptotic (cleaved Caspase 3) cells in forelimb bud sections from E10.5 mutant and control embryos (**Fig. S5**). None of these were detectably altered, suggesting that perturbation of cell proliferation and apoptosis were unlikely to be major contributors to the alterations observed in these embryos. Together with the finding that limb skeletal element differentiation did not appear to be significantly impaired or delayed in mutant embryos (**Fig. S2**), these data suggest that differences in cell proliferation and apoptosis in the limb bud are unlikely to account for most later phenotypic abnormalities observed in mutant limbs.

**Figure 4.**
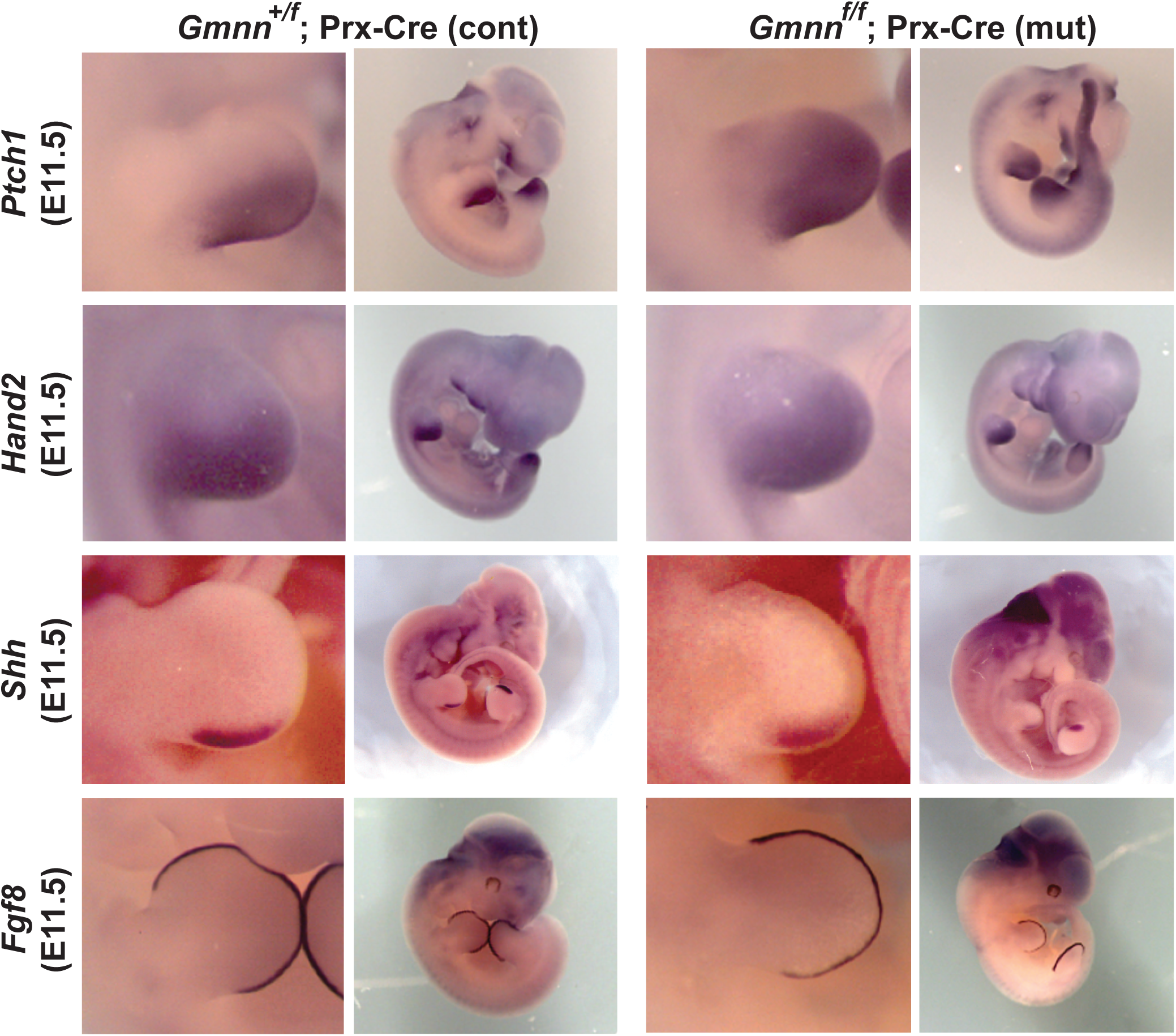
Forelimb buds of *Gmnn^f/f^*; Prx-Cre mutant versus *Gmnn^+/f^*; Prx-Cre control embryos exhibit similar expression of other classes of markers. Somite-matched pairs of mutant and control embryos from the same timed pregnancy were used for WISH with probes specific for the genes shown. A summary of findings, including number of biological replicates for each WISH probe, is provided in Table S3A.

To further examine how gene expression was altered during limb development in *Gmnn^f/f^*; Prx-Cre embryos, we also collected somite-matched forelimb buds from pairs of E10.5 mutant and control embryos, with each pair obtained from the same timed pregnancy, and performed RNA-seq analysis to define differentially expressed genes (DEGs) (see Methods). The majority of DEGs exhibited lower expression in *Gmnn^f/f^*; Prx-Cre forelimb bud samples, relative to *Gmnn^+/f^*; Prx-Cre control-derived samples (**Fig. S6A, Table S4**). We assessed the potential biological functions of these DEGs by identifying terms and networks associated with all DEGs, or only with up- or down-regulated DEGs (**Table S4**). Among all DEGs, many top terms related to general developmental processes, including transcription, growth, and differentiation, as well as morphology of bone, connective tissue, and muscle, which are tissue derivatives found in the later limb (**Fig. 5A**). These terms include DEGs with known roles in limb development, including *Sox9*, *Sox11*, *Kdm6b, Notch2*, and *Meis1* (**Fig. 5B-D**). As most of these genes encode transcriptional regulators, they were also associated with the ‘transcription’ term, which also included *Lbx1, and Kmt2d* (**Fig. S6B**). Networks built from these DEGs are associated with development, and cell survival, growth, and proliferation (**Fig. S7A-B**). More specifically, we assessed only networks downregulated in the mutant limb bud compared to the control, as the majority of DEGs fell into this category. Top networks were associated with embryonic development, as well as developmental and connective tissue disorders (**Fig. S7C-D**). This suggests that the aberrant patterning of the mutant forelimb bud may alter expression of genes important for development of limb elements, at least partially contributing to the phenotypes observed.

**Figure 5.**
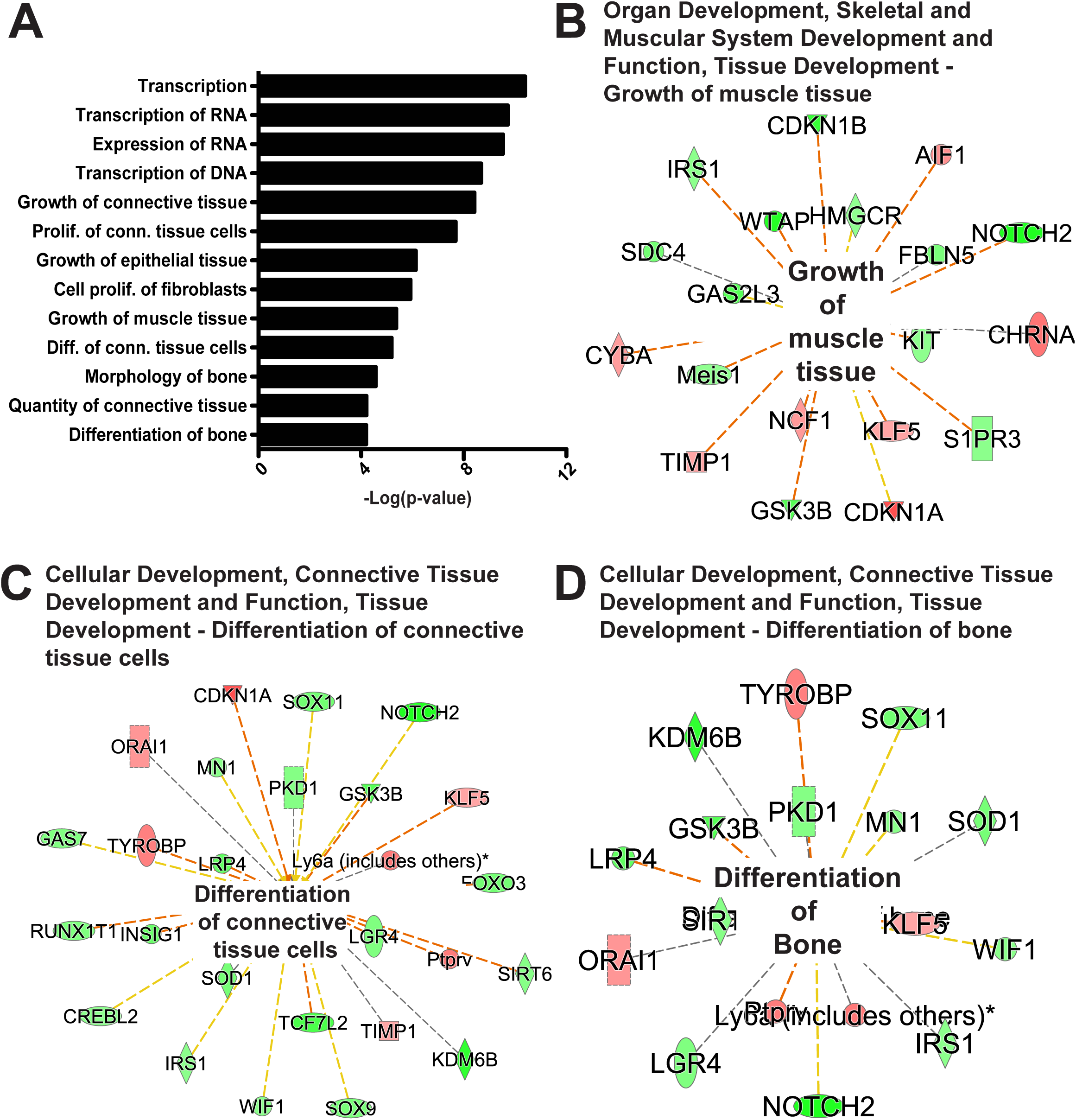
Differences in gene expression in *Gmnn^f/f^*; Prx-Cre mutant versus *Gmnn^+/f^*; Prx-Cre control forelimb buds. Forelimb buds were dissected from six pairs of somite-matched mutant and control E10.5 embryos (with each pair obtained from the same timed pregnancy) and subjected to RNA-seq analysis, as described in the methods. (A) Selected top class- and function-associated terms associated with DEGs, as identified by Ingenuity Pathway Analysis (IPA). (B-D) Genes within a selection of these terms are shown. Within each term, red symbols indicate upregulated genes and green symbols indicate downregulated genes, where the color intensity represents relative degree of differential expression.

### Ectopic ZPA formation in hindlimb buds of *Gmnn^f/f^*; *Dermo^Cre^* embryos

The *Gmnn^f/f^; Dermo^Cre^* mutant hindlimb buds exhibited polydactyly. We hypothesized that this might involve ectopic, anterior expression of key posterior patterning genes to generate an ectopic ZPA, as is seen in other models involving polydactyly (Lopez-Rios, 2016; Tickle and Towers, 2017). Since the *Gmnn^f/f^*; Prx-Cre forelimb buds exhibited altered patterns of *Hox* gene expression, we also hypothesized that 5’ *Hox* gene misexpression could potentially be involved in this mutant phenotype. Analysis of gene expression by WISH demonstrated that the E11.5 hindlimb buds of *Gmnn^f/f^; Dermo^Cre^* mutant embryos exhibited, with incomplete penetrance, ectopic anterior domains of expression of both *Shh* and *Ptch1*, a key target of the SHH signaling pathway, and also ectopically expressed *Hoxd13* in the same anterior location (**Fig. 6A-C**; **Table S3**). During limb bud patterning, a cleaved form of GLI3 transcription factor that acts as a transcriptional repressor (GLI3R) prevents anterior formation and expansion of ZPA activity. Therefore, we collected somite-matched pairs of E11.5 embryos with the *Gmnn^f/f^; Dermo^Cre^* (mutant) and *Gmnn^+/f^; Dermo^Cre^* (control) genotypes from single timed pregnancies and generated lysates from the fore- and hindlimb buds. These limb bud lysates were analyzed for levels of both the full-length transcriptional activator form of GLI3 (GLI3A) and for GLI3R, using an antibody that recognizes both forms of the protein. GLI3A levels were unchanged in both the mutant fore- and hindlimb-buds, while GLI3R levels were unchanged in the mutant forelimb bud, but were reduced in the mutant hindlimb bud (**Fig. 6D**). This finding is congruent with the finding of ectopic ZPA-like activity in the anterior region of these embryos, as this additional SHH signaling would be expected to antagonize cleavage of GLI3 to GLI3R (Lopez-Rios, 2016; Tickle and Towers, 2017).

**Figure 6.**
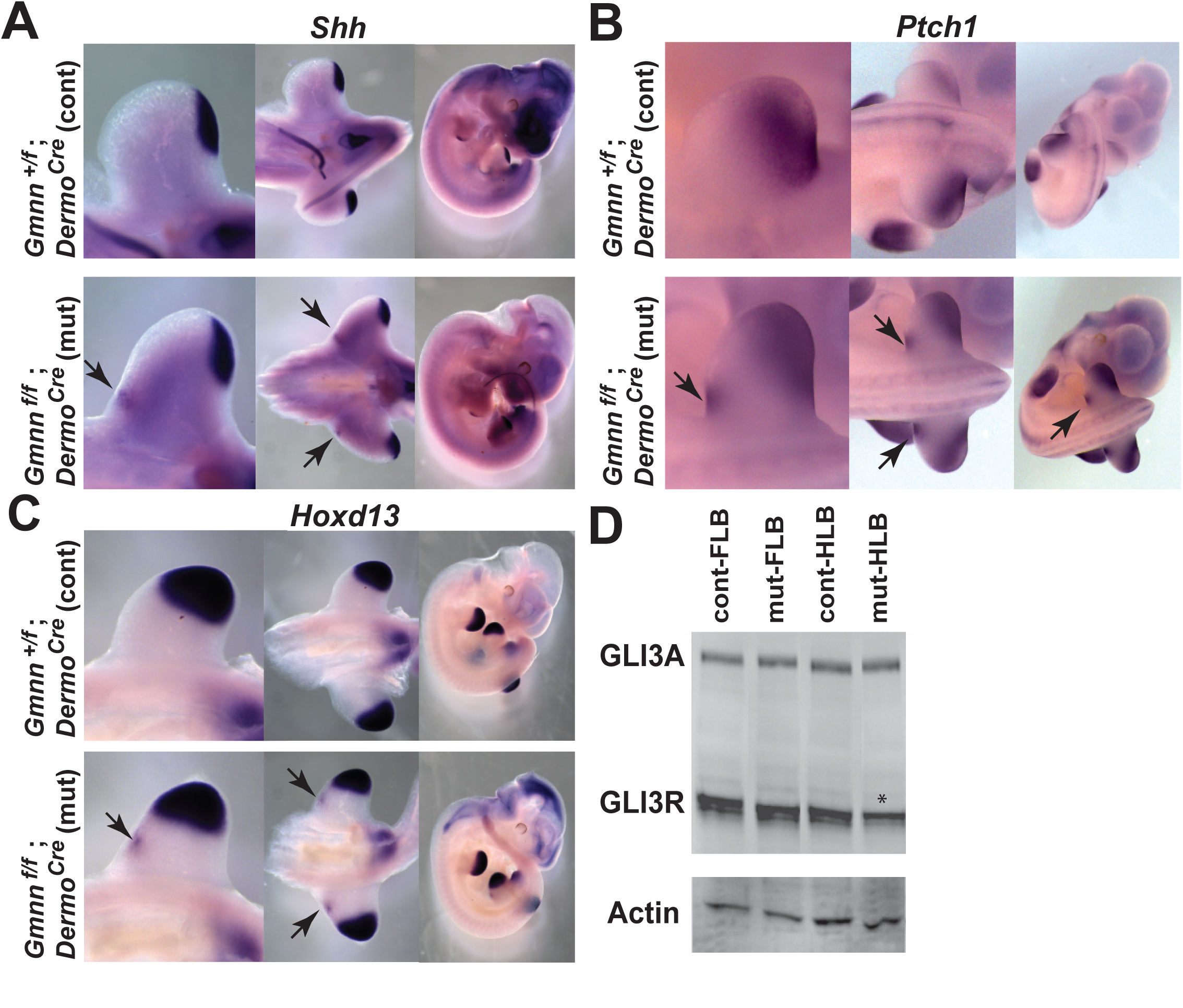
Ectopic expression of *Shh*, *Ptch1*, and *Hoxd13*, and reduced GLI3R levels in hindlimb buds of *Gmnn^f/f^; Dermo^Cre^* mutant embryos. Pairs of *Gmnn^+/f^; Dermo^Cre^* control and *Gmnn^f/f^; Dermo^Cre^* mutant somite-matched E11.5 embryos obtained from the same timed pregnancy were used for (A-C) WISH or (D) isolation of fore- and hindlimb bud tissue lysates for Western blotting. (A-C) Hindlimb bud, posterior trunk, and whole embryo side views are shown, with arrows marking ectopic expression in the anterior hindlimb bud of the mutant. (D) Immunoblotting of hindlimb buds with a GLI3 antibody detects the full length activating (GLI3A) and cleaved repressor (GLI3R) forms, the latter of which is reduced in the mutant hindlimb bud (* highlights reduced expression; n=3 biological replicate experiments). A summary of findings, including number of replicates for each WISH probe, is provided in Table S3B.

We also examined the expression of other markers by WISH in *Gmnn^f/f^; Dermo^Cre^* mutant and *Gmnn^+/f^; Dermo^Cre^* control embryos, again using somite-matched pairs of embryos of each genotype from the same timed pregnancy for analysis. Expression of the posterior marker, *Hand2*, and the AER marker, *Fgf8*, were similar in both control and mutant embryos (**Fig. 7**). Other than the ectopic anterior domain of *Hoxd13* expression observed above, expression of most other 5’ *Hox* genes examined (*Hoxd10, Hoxd11, Hoxd12, Hoxa13*) appeared similar in control and mutant embryos at E11.5 (**Fig. 7**). However, while *Hoxd12* expression did not differ between mutant and control embryos at E11.5, by E12.5, its expression remained excluded from the anterior portion of the hindlimb bud in control embryos, but spread ectopically throughout the anterior portion of the distal hindlimb bud in mutant embryos (**Fig. 7**, **arrows**). As only *Hox13* paralogs are expressed in this anterior portion of the forming autopod during normal limb development, the finding that *Hoxd12* expression had expanded into this region in the mutant hindlimb autopod reflects an alteration of anterior-posterior patterning and could contribute to the hindlimb polydactyly observed at later stages.

**Figure 7.**
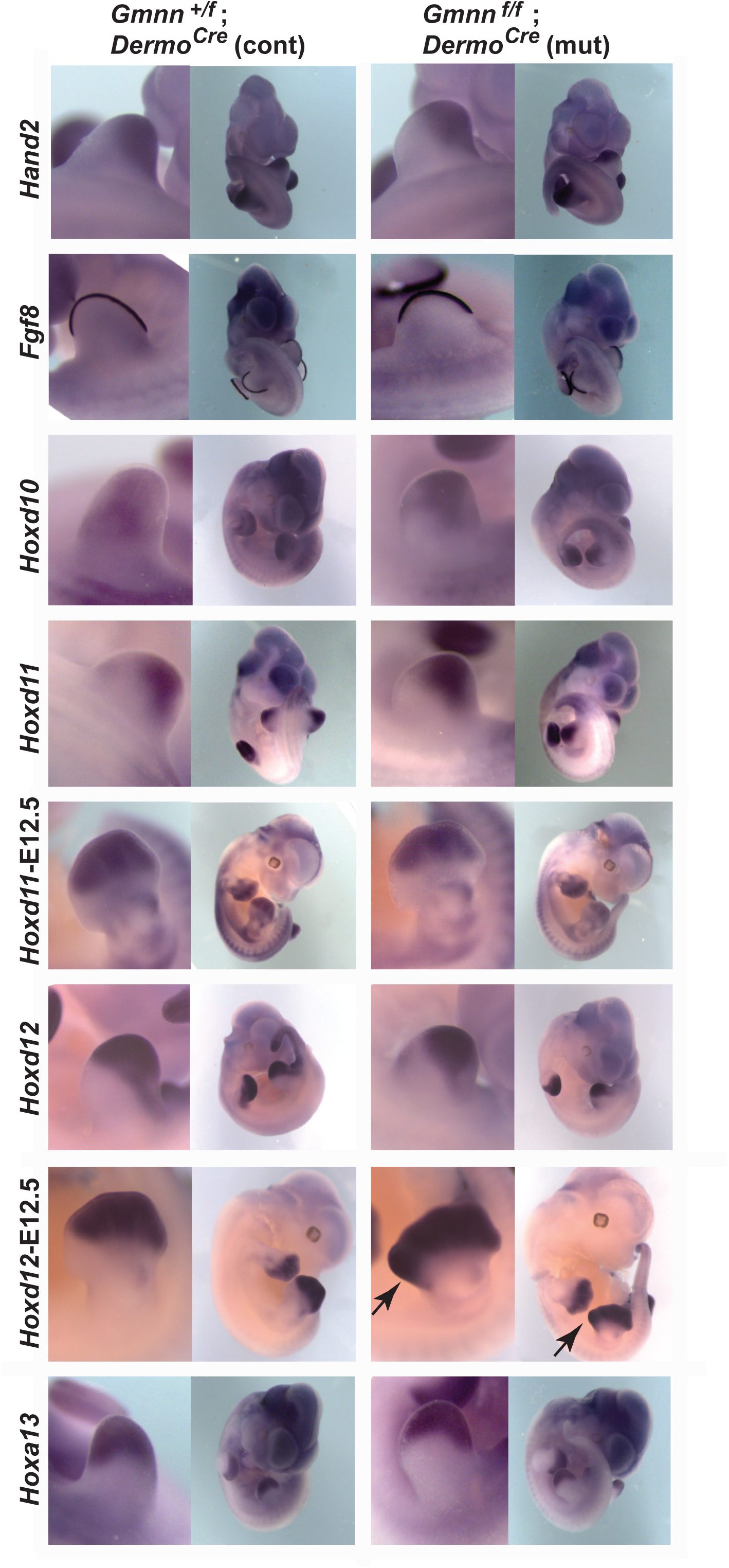
Hindlimb buds in *Gmnn^f/f^; Dermo^Cre^* mutant and *Gmnn^+/f^*; *Dermo^Cre^* control embryos exhibit similar expression of *Hand2*, *Fgf8*, and most 5’ *Hox* genes. Pairs of somite-matched mutant and control embryos from the same timed pregnancy were analyzed at E11.5 (or at E12.5, where indicated) by in WISH with probes specific for the genes indicated. An arrow marks expanded *Hoxd12* expression into the anterior or the autopod in the mutant at E12.5. A summary of findings, including number of replicates for each WISH probe, is provided in Table S3B.

## Discussion

While *Geminin* (*Gmnn*) plays several roles in patterning the primary embryonic body axis, a role in limb development had not previously been suggested. Here, we used conditional *Gmnn* knockout with a Prx-Cre driver to define a requirement for *Gmnn* for proper development of forelimb skeletal elements. During embryonic patterning of *Gmnn*-deficient forelimb buds, 5’ *Hox* gene expression expanded into more proximal and anterior regions of the limb bud, while mutant animals later exhibited loss or reduction of stylopod- and zeugopod-derived skeletal elements. Conditional *Gmnn* knockout with a *Dermo^Cre^* driver instead resulted in hindlimb polydactyly and in similar, but less severe, reductions of stylopod- and zeugopod-derived hindlimb skeletal elements in some mutant animals. The developing hindlimb buds in these mutant embryos formed an ectopic anterior ZPA, which misexpressed *Hoxd13, Shh*, and *Ptch1*, while GLI3 repressor levels were also reduced in these mutant limb buds and late expression of *Hoxd12* expanded anteriorly. Together, this work identified several new requirements for *Gmnn* activity for normal patterning of the vertebrate limb.

### Comparison of conditional models of Geminin deficiency in mesodermal derivatives

The observations that *Gmnn^f/f^;* Prx-Cre mutant animals had disrupted skeletal development in the forelimbs but not hindlimbs, while *Gmnn^f/f^; Dermo^Cre^* animals had normal forelimb but altered hindlimb development, suggest a time period during which limb development may be particularly sensitive to *Gmnn* deficiency. At E9.5, *Dermo^Cre^* is expressed in the hindlimb ridge but not in the forelimb bud, while, by E10.5 it is expressed in both the fore- and hindlimb buds. Induction of *Gmnn* deficiency with this driver resulted only in hindlimb defects, suggesting that forelimb development is most sensitive to *Gmnn* loss prior to E10.5. Congruent with this, Prx-Cre is already expressed in the FLB from E9.5, and excision using this driver altered forelimb development, while Prx-Cre expression is only weakly detected in the hindlimb bud by E10.5, and *Gmnn^f/f^;* Prx-Cre animals exhibited normal hindlimb development. As early aspects of limb bud patterning occur approximately a half day later in the hindlimb versus forelimb buds (Lopez-Rios, 2016), the differential effects seen in these models suggest a window of sensitivity to *Gmnn* deficiency corresponding to early aspects of limb bud outgrowth and patterning (e.g. from E9.5-10.5 in the forelimb).

Interestingly, *Gmnn* excision with these Prx-Cre and *Dermo^Cre^* drivers did not result in axial skeletal phenotypes, despite the fact that both genes are expressed in some mesodermal tissues of the primary body axis from E9.5 (Logan et al., 2002; Yu et al., 2003). By contrast, earlier excision in axial mesoderm with the T-Cre driver, which is expressed in gastrula mesoderm from E7.5 (Perantoni et al., 2005) resulted in embryonic lethality by E9.5. Embryos isolated at these stages exhibited loss or gross reduction of all caudal tissues, while these caudal tissues displayed elevated apoptosis by E8.5. The cause of this apoptosis is unclear. In the head and trunk of *Gmnn^f/f^*; T-Cre; mutant embryos at E8.5, immunohistochemical analysis revealed well patterned trunk tissues expressing mesodermal, endodermal, and neural markers, suggesting that apoptosis is not a consequence of grossly disrupted specification and patterning of these axial tissues. As GMNN acts within one of several, often redundantly functioning mechanisms to maintain the fidelity of DNA replication (Nishitani et al., 2004; Saxena and Dutta, 2005; Sugimoto et al., 2004; Zhu and Depamphilis, 2009), apoptosis could be triggered by DNA re-replication and subsequent genome abnormalities in some cells. However, we would not expect the effects of such cell cycle-related perturbation to be restricted to the caudal end of the embryo as is seen in this model. Alternatively, apoptosis could potentially be triggered by impaired mesoderm migration through the primitive streak at the caudal end of the embryo during gastrulation. *Gmnn* knockdown in the early mouse embryo by shRNA injection at E6.0 was previously reported to disrupt the morphogenetic movements of gastrulation and to impair anterior-posterior extension, resulting in embryonic lethality around E9.5, with the fraction of embryos that survived to this stage failing to complete turning (Emmett and O’Shea, 2012). The phenotype described here using T-Cre-mediated *Gmnn* excision is reminiscent of these findings and is congruent with this previously reported requirement for *Gmnn* activity in gastrula mesoderm (Emmett and O’Shea, 2012). Likewise, altered DNA replication is unlikely to be a major contributor to the limb-specific phenotypes observed in the Prx-Cre and *Dermo^Cre^* models, as markers of apoptosis and proliferation were unaffected, while 5’ *Hox* gene expression territories were expanded, in the limb bud of Prx-Cre mutant embryos at this time.

### Reduction and loss of forelimb skeletal elements in the *Gmnn^f/f^*; Prx-Cre model

We explored potential molecular contributors to the limb phenotypes seen in our Prx-Cre and *Dermo^Cre^* models during limb bud patterning and outgrowth. During other developmental processes, GMNN physically and/or functionally cooperates with Polycomb (PcG)-mediated repression to maintain normal spatial and temporal expression of *Hox* genes (Karamitros et al., 2014; Luo et al., 2004). GMNN also cooperates with PcG activity to restrain developmental gene expression during mesendoderm specification (Caronna et al., 2013; Lim et al., 2011). Therefore, we hypothesize that the limb phenotypes observed here could reflect a parallel requirement for *Gmnn* to maintain proper spatial and temporal patterns of *Hox* gene expression in the developing limb buds. Indeed, in *Gmnn^f/f^*; Prx-Cre mutant forelimb buds, expression of 5’ *HoxD* cluster genes was expanded, with all *Hox* genes examined exhibiting ectopic expression in proximal and anterior limb bud domains from which they were restricted in control embryos (schematized in **Fig. 8A**). This suggests that, during early limb bud formation and outgrowth, mutant embryos had diminished capacity to restrict the limb bud regions in which *HoxD* cluster gene expression was activated and maintained.

**Figure 8.**
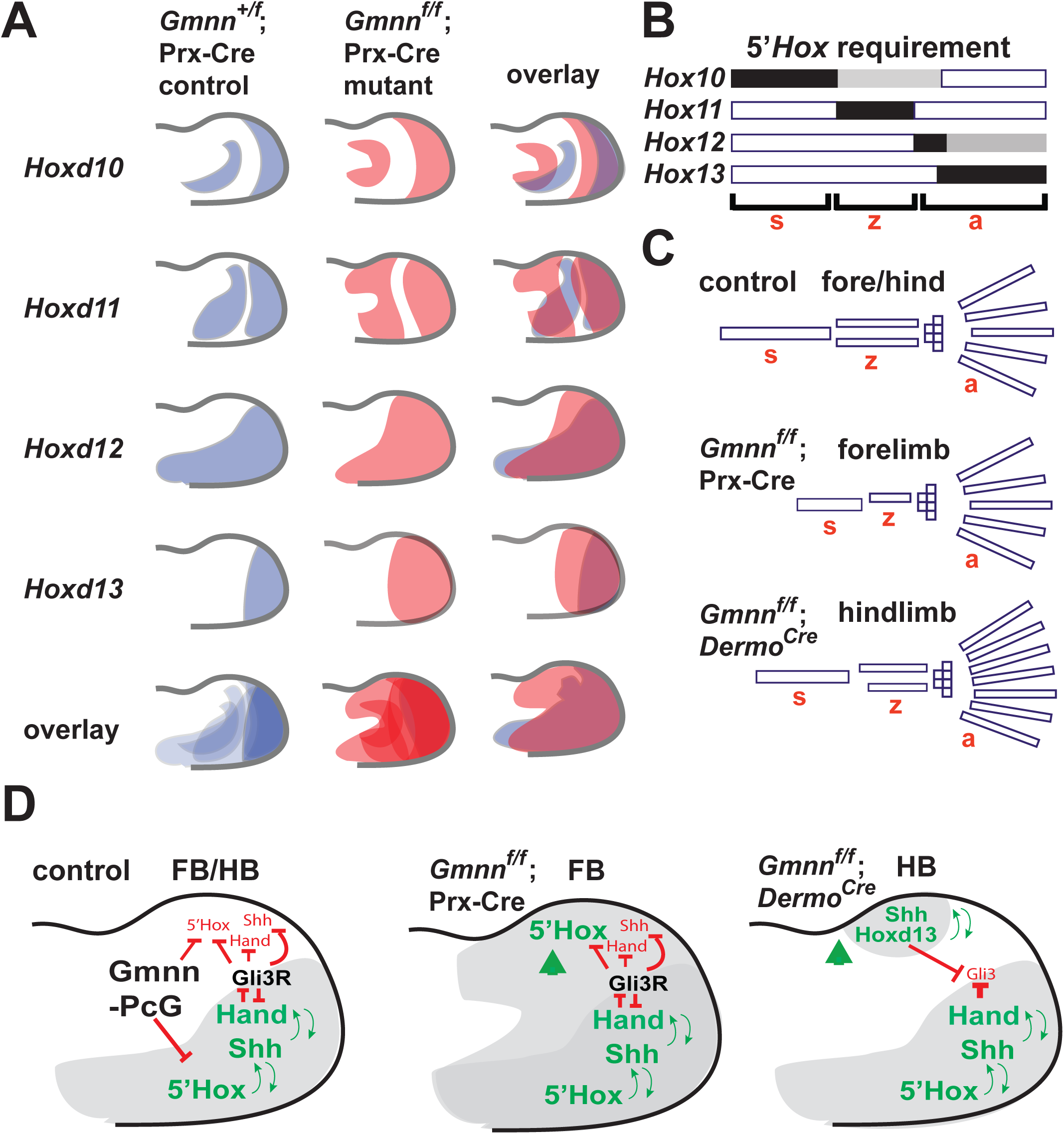
Modeling Geminin requirements for limb development. (A) 5’ *HoxD* cluster genes are expressed in an overlapping pattern, with *Hoxd13* exhibiting the most distally restricted expression. *HoxD* expression domains are expanded in the *Gmnn^f/f^*; Prx-Cre mutant forelimb buds by comparison with *Gmnn^+/f^*; Prx-Cre controls, with the expansion visible in the overlay of control versus mutant expression domains. (B) Different 5’ *Hox* paralogs are required (shown in black, with a lesser requirement indicated in gray) for formation of each skeletal element, with *Hox10*, *Hox11* and *Hox13* paralogs controlling formation of stylopod (s), zeugopod (z), and autopod (a) elements, respectively. (C) Schematics of bone elements corresponding to the domains of *Hox* requirement above, with *Gmnn^f/f^*; Prx-Cre mutant embryos exhibiting shorter or missing limb skeletal elements, while *Gmnn^f/f^*; *Dermo^Cre^* mutant embryos exhibit some skeletal element shortening and polydactyly. (D) Model for *Gmnn* requirements in patterning the limb bud. GMNN may act cooperatively with PcG to restrain 5’ *Hox* gene expression in the developing limb bud, modulating *Hox* expression domains. In the *Gmnn* deficient forelimb bud, expansion of 5’ *Hox* expression into more proximal and anterior limb bud regions may disrupt regional specification of limb elements while, in the hindlimb bud, *Gmnn* deficiency results in the formation of a *Hoxd13*- and *Shh*-expressing ectopic, anterior ZPA signaling center, altering later limb patterning. FB – foreblimb bud, HB-hindlimb bud.

5’ *Hox* cluster gene regulation must be finely tuned to generate a ‘nested’ pattern of expression corresponding to the location of each gene in the cluster, with the most 5’ gene in the cluster, *Hoxd13*, exhibiting the most distally restricted expression (Andrey and Duboule, 2014; Montavon and Duboule, 2013; Montavon and Soshnikova, 2014; Neijts and Deschamps, 2017; Noordermeer and Duboule, 2013). Expressing the incorrect complement or dose of *HoxD* genes in more anterior and proximal regions of the limb bud, as is seen here, would be expected to alter multiple aspects of later limb patterning, as *Hox10, Hox11*, and *Hox13* paralogs play essential roles in regulating the development of stylopod-, zeugopod-, and autopod-derived structures, respectively (**Fig. 8B**) (Davis et al., 1995; Fromental-Ramain et al., 1996b; Wellik and Capecchi, 2003). Interestingly, while nearly all mutant limbs analyzed exhibited loss or reduction of stylopod and/or zeugopod-derived skeletal elements, as schematized in **Fig. 8C**, the specific nature of the abnormalities varied. Some limbs contained all skeletal elements, but exhibited moderate to severe shortening of one or all elements. Other mutant limbs exhibited loss of either the stylopod- and/or one or both zeugopod-derived skeletal elements. This phenotypic variability may reflect a role for *Gmnn* in fine tuning 5’ *HoxD* expression during limb bud patterning and outgrowth.

Temporal and spatial control of *Hox* expression in the developing limb bud is under complex control (Andrey and Duboule, 2014; Montavon and Duboule, 2013; Montavon and Soshnikova, 2014; Neijts and Deschamps, 2017; Noordermeer and Duboule, 2013), with the exact complement of *Hox* gene expression differing even among cells within the same region of the limb bud (Fabre et al., 2018). Polycomb is a central modulator of 5’ *Hox* spatial and temporal expression in the developing limb bud. Removal of PcG-mediated H3K27me3 from 5’ *Hox* genes in the most posterior mesenchyme is required for their activation (Montavon and Duboule, 2013; Soshnikova and Duboule, 2009; Williamson et al., 2012), which also involves large scale chromatin reorganization, including chromatin decompaction and long-range interactions with distant enhancers located telomeric or centromeric to the gene cluster (Andrey et al., 2013; Kmita et al., 2002; Montavon et al., 2011; Tschopp et al., 2009); reviewed in (Andrey and Duboule, 2014; Montavon and Duboule, 2013; Montavon and Soshnikova, 2014; Neijts and Deschamps, 2017; Noordermeer and Duboule, 2013). As these highly orchestrated regulatory processes coordinate spatial and temporal activation of *Hox* gene expression to generate a precisely nested pattern in the forming limb bud, their disruption is likely to be a significant contributor to the alterations of limb development observed in these models, and could involve GMNN’s capacity for cooperativity with Polycomb to restrict *Hox* expression (Karamitros et al., 2014; Luo et al., 2004).

In support of this model, schematized in **Fig. 8D**, the phenotypes observed in these *Gmnn* conditional knockout models are reminiscent of other mutations that result in either loss or gain of *Hox* gene expression. For example, loss of function of *Hoxa10, Hoxc10*, and *Hoxd10* or of both *Hoxa11* and *Hoxd11* in mouse models severely affects the development of stylopod- or zeugopod-derived skeletal elements, respectively (Davis et al., 1995; Wellik and Capecchi, 2003). Reduction of specific limb skeletal elements (e.g. the zeugopod) without alterations of autopod development, as is seen in the *Gmnn^f/f^;* Prx-Cre model, also resembles limb defects seen in models involving *Hox* misexpression. For example, the semi-dominant mouse mutation Ulnaless results in severe reduction of zeugopod-derived skeletal elements in both the forelimb and hindlimb, defects evident by E14.5 (Peichel et al., 1997). During earlier patterning, Ulnaless limb buds exhibit ectopic *Hoxd12* and *Hoxd13* expression, which expands from the distal into the proximal limb bud. Such *Hoxd12/13* misexpression appears to interfere with the ability of *Hox* group 11 gene activity to elaborate the zeugopod skeletal element, contributing to the Ulnaless phenotype (Peichel et al., 1997). Conditional knockout of the PcG gene *Ezh2* likewise triggers expanded and ectopic *Hox* 5’ gene expression (Wyngaarden et al., 2011). Reminiscent of the *Gmnn^f/f^; Dermo^Cre^* phenotype observed here, the limb buds of these *Ezh2* conditional knockout embryos similarly exhibit ectopic anterior expression of *Ptch1* and reduced expression of *Gli3*, while the mutant animals later exhibit shortening of all three primary proximo-distal limb segments (stylopod, zeugopod, and autopod) as well as anterior-posterior patterning anomalies involving the autopod (Wyngaarden et al., 2011).

In the *Gmnn^f/f^*; Prx-Cre model, the autopod may be patterned normally due to the posterior dominance of *Hoxd13* activity over the activities of the other 5’ *HoxD* genes. These differential effects could also be influenced by temporal differences in regional specification of the stylopod/zeugopod versus the autopod, as these are controlled by earlier versus later waves of *Hox* transcriptional regulation, which are under distinct regulatory controls (Andrey and Duboule, 2014; Andrey et al., 2013; Montavon and Duboule, 2013; Montavon and Soshnikova, 2014; Neijts and Deschamps, 2017; Noordermeer and Duboule, 2013). It should also be noted that the 5’ *Hox* genes contribute to several aspects of later limb development, such as regulation of chondrogenic differentiation and maturation to generate limb skeletal elements (Boulet, 2003; Swinehart et al., 2013). In the *Gmnn^f/f^*; Prx-Cre model, persistence of an incorrect level or aberrant combinations of 5’ *HoxD* gene expression in regions of the limb bud could impair these later processes, in addition to altering early limb bud patterning. However, loss and reduction of limb skeletal elements was already apparent in mutant embryos by E14.5, and differentiation of the skeletal elements that were present did not appear appreciably impaired when examined at later stages (**Fig. S2**). Misexpression of the *Hox* 5’ genes was already visible during earlier patterning of the limb bud at E11.5, suggesting that this may be a major contributor to the skeletal defects observed at later stages.

To explore the molecular basis of the limb phenotypes seen in the *Gmnn^f/f^*; Prx-Cre model, beyond analyzing expression of key limb developmental regulatory genes by WISH, we performed RNA-seq on pairs of somite-matched mutant and control E10.5 embryos, each derived from the same timed pregnancy. Even at this early stage of forelimb development, some developmental regulatory genes were differentially expressed. Many of these genes encode proteins with roles in regulating the development of derivatives of limb bud mesenchyme, such as bone, muscle, and connective tissue, while mutations in some of these genes have been connected to limb-associated phenotypes in mice and/or humans. For example *Sox9* and *Sox11* are involved in cartilage condensation (Bhattaram et al., 2014; Healy et al., 1999), *Meis1* plays a role in specification of stylopod-derived structures (Ye et al., 2016), *Notch2* regulates limb mesenchyme development during interdigital apoptosis (Pan et al., 2005), *Cux2* modulates the proliferation to differentiation transition in the developing limb (Tavares et al., 2000), and *Lbx1* controls limb muscle precursor migration into the limb (Gross et al., 2000; Schafer and Braun, 1999). Mice with mutations in a number of the DEGs identified here exhibit abnormal limb development and morphology, including *Kdm6b* (Zhang et al., 2015), *Lrp4* (Simon-Chazottes et al., 2006), *Slc39a1* (Dufner-Beattie et al., 2006), *Lbx1* (Gross et al., 2000), and *Sox9* (Bi et al., 2001; Yap et al., 2011). Mutations in some of these DEGs also cause human syndromes involving limb and axial skeletal developmental abnormalities including *Sox9* (campomelic dysplasia; osteochondrodysplasia; bowing of tibia/femur; OMIM608160; (Unger et al., 1993)), *Lbr* (Greenberg skeletal dysplasia; OMIM600024), *Lrp4* (Sclerosteosis; bone dysplasia and syndactyly of fingers and toes and Cenani-Lenz Syndactyly Syndrome-shortening of the radius and ulna; OMIM604270), and *Kmt2d* (Kabuki syndrome; short stature and skeletal anomalies; OMIM602113). Interestingly, while most transcriptional targets of 5’*Hox* cluster genes in the developing limb are not known, a comparative dataset for HOXD13 targets was defined by ChIP in a mesenchymal cell line and includes a number of these DEGs, including *Meis1*, *Kit, Epha7, Rab3c, Btg2, Ccng1, Pcdhgb6, Abhd2, Pcdhga7, Pcdhga5,* and *Eif4g2* (Salsi et al., 2008). This suggests that *Gmnn* deficiency perturbs gene regulatory networks needed for patterning of the limb bud and controlling the specification and differentiation of limb bud-derived tissues.

### Hindlimb polydactyly and skeletal element shortening in the *Gmnn^f/f^*; *Dermo^Cre^* model

The phenotype of *Gmnn^f/f^; Dermo^Cre^* embryos differed from that seen in *Gmnn^f/f^;* Prx-Cre embryos, in that the hind-rather than forelimbs were affected, and the predominant phenotype was autopod polydactyly, with most mutant limbs having 6 or 7 digits (92% of limbs). This comparison is schematized in **Fig. 8C**. Polydactyly generally involves production of an ectopic, anterior ZPA and, indeed, *Hoxd13*, the HOXD13 target *Shh*, and the SHH target gene *Ptch1* were all expressed in an ectopic anterior domain in mutant hindlimb buds. Ectopic expression of these genes was observed with incomplete penetrance, likely due to the limited sensitivity of in situ hybridization and the ectopic expression of these transcripts in a relatively small number of cells in a spatially restricted domain of the developing limb bud. In other mouse models, such ectopic anterior *Shh* expression confers ZPA activity and likewise drives polydactyly, suggesting that this molecular misregulation is likely to underlie the polydactyly phenotype seen here (Lopez-Rios, 2016; Tickle and Towers, 2017).

It is not clear how *Gmnn* deficiency leads to activation of this ectopic anterior domain of *Shh* expression in the developing hindlimb bud. We hypothesize that this may also involve transient misexpression of 5’ *HoxD* cluster genes in anterior limb bud cells during limb bud pre-patterning. Expression of the 5’ *HoxD* genes is normally restricted to the posterior of the limb bud during a pre-patterning stage; this occurs prior to and independent of *Shh* expression and subsequently promotes its expression, in part through direct binding of both HOXD13 and HAND2 to a long-range *Shh* enhancer that promotes its expression in the limb bud (denoted the ‘zone of polarizing activity regulatory sequence’ or ZRS). During limb bud pre-patterning, the cleaved repressor form of GLI3 (GLI3R) blocks anterior expansion of 5’ *Hox* gene expression, while GLI3R also directly antagonizes binding of HAND2 and HOXD13 binding to and activation of the *Shh* ZRS, and long-range looping of this enhancer to the *Shh* promoter (Galli et al., 2010; Hill, 2007; Lettice et al., 2003; Lopez-Rios, 2016; Sagai et al., 2005; Tickle and Towers, 2017). HAND2 activity is posteriorly restricted and excluded from the anterior limb bud by mutual antagonism with GLI3 at this time. Once *Shh* expression is initiated, *Shh* and the 5’ *HoxD* genes establish positive feedback loops to promote each other’s posterior expression and to antagonize GLI3R production (Lopez-Rios, 2016; Tickle and Towers, 2017). In the mutant, these feedback loops may maintain *Shh* expression in both the endogenous and ectopic ZPA domains, increasing antagonism of GLI3R production (schematized in **Fig. 8D**). In *Gmnn^f/f^*; *Dermo^Cre^* embryos, an ectopic ZPA feedback loop appears to have been generated, marked by ectopic *Hoxd13, Shh*, and *Ptch1* expression, and corresponding with reduced GLI3R levels in mutant hindlimb buds, potentially involving antagonism of GLI3R production by both the endogenous and ectopic ZPAs. In addition to restricting 5’ *HoxD* gene expression, GMNN can also interact directly with a number of the HOX transcription factors, including HOXD13, antagonizing their transcriptional activity (Salsi et al., 2009; Zhou et al., 2012). Therefore, if GMNN and HOXD13 proteins interact in the limb bud *in vivo*, this interaction could potentially constrain the ability of HOXD13 to bind the *Shh* ZRS, such that GMNN loss could increase the capacity of HOXD13 to activate this enhancer.

Although the *Gmnn^f/f^; Dermo^Cre^* phenotype is distinct from the *Gmnn^f/f^*; Prx-Cre phenotype, reduced capacity to restrict or fine-tune expression of 5’ *Hox* genes may contribute to both phenotypes, as schematized by the altered 5’ *Hox* expression present in both *Gmnn* mutant limb bud models in **Fig. 8D**. In bone preparations from the *Gmnn^f/f^; Dermo^Cre^* animals, we did not observe loss of any hindlimb skeletal elements; however, 59% of the adult hindlimbs exhibited shortening and curvature of both stylopod- and zeugopod skeletal elements, reminiscent of a less severe form of the phenotype seen in the *Gmnn^f/f^*; Prx-Cre model. Unlike results for the *Gmnn^f/f^*; Prx-Cre model, in the *Gmnn^f/f^; Dermo^Cre^* limb buds we did not observe overt expansion of the expression of most 5’ *Hox* genes into more anterior or proximal regions of the limb bud. However, we did observe late anterior misexpression of *Hoxd12* throughout the anterior portion of the autopod, in a territory from which its expression would normally be excluded. Such *Hox* misexpression may contribute to the altered anterior-posterior patterning of the limb bud and suggests that misexpression of 5’ *Hox* genes, while not as severe as was seen for the *Gmnn^f/f^*; Prx-Cre limb buds, may also contribute to abnormalities in formation of stylopod- and zeugopod-derived skeletal elements that are seen in both models.

Here we showed that *Gmnn* loss resulted in dramatic alterations of the limb patterning program, altering 5’ *Hox* gene expression. This ultimately resulted in different limb developmental phenotypes in each conditional model, highlighting the dependence of this process on the precise control of 5’ *Hox* gene expression. Expression of the *Hox* gene clusters is subject to multiple layers of regulatory controls, each involving complex activities of both local and long-range enhancers, and also involving orchestrated PcG complex activity (Andrey and Duboule, 2014; Montavon and Duboule, 2013; Montavon and Soshnikova, 2014; Neijts and Deschamps, 2017; Noordermeer and Duboule, 2013). Given the heterogeneous and dynamic nature of this regulation, including variations in gene expression across the limb bud at the level of the single cell (Fabre et al., 2018), improvements in technology to couple spatial resolution with single-cell interrogation of how these regulatory controls operate should allow us to more precisely elucidate the regulatory mechanisms underlying limb patterning.

## Methods

### Mouse strains and husbandry

*Dermo^Cre^* [Twist2^tm1(cre)Dor^](Yu et al., 2003) was obtained from Dr. David Ornitz. Prx1-Cre [Tg(Prrx1-cre)1Cjt], generated by Dr. Cliff Tabin, was obtained from Jackson laboratory (Logan et al., 2002). T-Cre [Tg(T-cre)1Lwd] was obtained from Dr. Mark Lewandoski (Perantoni et al., 2005). T-Cre expresses Cre recombinase from a 500 bp *brachyury* (*T/Bry*) promoter in mesodermal lineages. At E6.5, no T-Cre expression is detected, while at E7.5 recombination occurs in the primitive streak and migrating mesoderm but not the node. By E8.5, Cre recombination occurs in paraxial, intermediate, and lateral mesoderm, and by E9.0 most mesodermal lineages express Cre (Perantoni et al., 2005). Geminin conditional knockout mice (CSD24729) were purchased from the Knockout Mouse Project (KOMP). The targeted *Gmnn* allele contains a splice acceptor-βgeo-polyA sequence flanked by FRT sites (permitting excision by FLPe recombinase) inserted into intron 3 and loxP sites flanking exon 4 (permitting excision of exon 4 by Cre recombinase). Prior to FLPe excision of the splice acceptor-βgeo-polyA sequence, the targeted allele is predicted to be a null allele of *Gmnn*. Excision by FLPe recombinase results in a targeted (floxed) allele that generates a wild-type mRNA. This floxed allele can be converted to a null allele in the presence of Cre, which excises exon 4 (Patterson et al., 2014). For timed matings, noon of the day of plug discovery was designated E0.5. All animals and embryos analyzed were generated by crossing *Gmnn^f/f^* females with either *Gmnn^+/f^*; Prx-Cre or *Gmnn^+/f^*; *Dermo^Cre^* males. Each biological replicate analysis was performed in parallel using animals, or somite-matched pairs of embryos obtained from the same timed pregnancy with the *Gmnn^+/f^*; Cre (control) or *Gmnn^f/f^*; Cre (mutant) genotypes. Animal studies were conducted under protocols approved by the Washington University Institutional Animal Care and Use Committee (Protocol #: 20190021).

### Genotyping

DNA was extracted from embryonic membranes or tail tissue using the HotShot method for 15 minutes (Truett et al., 2000) and 2 µl DNA was used for PCR with Phusion polymerase (New England Biolabs). For Geminin and Cre PCRs, cycling conditions were: 98°C for 60 seconds followed by 33 cycles of (98°C 10 seconds, 65°C 10 seconds, 72°C 30 seconds). Primers for genotyping were previously described (Patterson et al., 2014).

### Skeletal preparations

Embryos were dissected free of membranes, de-skinned, and fixed for 24 hours in 95% ethanol, followed by 24 hours in acetone. Embryos were stained overnight in 70% ethanol/5% acetic acid/0.015% Alcian Blue/0.005% Alizarin Red. Maceration was performed in 1% potassium hydroxide/20% glycerol until embryos were clear, as previously described (Colvin et al., 1996). After further clearing in 50% and 80% glycerol, embryos were photographed on a Zeiss Stereo Discovery V12 microscope.

### LacZ staining and hematoxylin and eosin staining

LacZ staining of E9.5-12.5 embryos was performed as previously described (Nagy et al., 2007). Hematoxylin and eosin (H&E) staining of E14.5-18.5 embryonic limb sections was performed by the Washington University in St. Louis Department of Developmental Biology histology core as previously described (Cardiff et al., 2014), with images captured using a Hamamatsu NanoZoomer.

### Immunofluorescence

Immunofluorescence was performed on frozen tissue sections of E8.5 embryos after prior methods (Wang and Matise, 2013). Antibodies used were: goat anti-Brachyury (AF2085; R&D Systems), goat anti-FOXA2 (M-20, sc-6554; Santa Cruz), goat anti-SOX1 (AF3369; R&D Systems), rabbit anti-Ki67 (90584; Thermo), rabbit anti-phospho-Histone H3 (06-570; Millipore), and rabbit anti-cleaved Caspase-3 (9661; Cell Signaling). Immunofluorescence imaging was performed on a Zeiss Axio Imager Z1 at 10X or 20X magnification.

### Whole Mount in Situ Hybridization (WISH)

Embryos were dissected free from extraembryonic membranes and were fixed in 4% paraformaldehyde overnight at 4°C. In situ hybridization was performed as previously described (Wilkinson and Nieto, 1993). Whole mount images of embryos were taken using a Zeiss Stereo Discovery V12.

### Western Blotting

Pairs of somite matched *Gmnn^f/f^*; *Dermo^Cre^* (mutant) and *Gmnn^+/f^*; *Dermo^Cre^* (control) E11.5 embryos, each isolated from the same pregnant female, were used to isolate fore- and hindlimb buds. Limb buds were used for protein lysate preparation and western blotting with the 6F5 anti-GLI3 antibody was performed as previously described (Wen et al., 2010). The 6F5 monoclonal antibody was provided by Suzie J. Scales (Genentech Inc.).

### RNA-seq and Data Analysis

Six litter- and somite-matched pairs of *Gmnn^f/f^*; Prx-Cre (mutant) and *Gmnn^+/f^*; Prx-Cre (control) E10.5 embryos were used to prepare RNA. RNA was quantified with the NanoDrop ND-1000 spectrophotometer (Thermo Scientific), and the quality of total RNA was further confirmed using the Agilent Bioanalyzer 2100. Samples were rejected if the RNA integrity number was below 8. RNA-seq library preparation (Ribo-zero) and Illumina HiSeq3000 sequencing were performed by the Genome Technology Access Center at Washington University. RNA-seq reads were aligned to the mouse genome mm10 with STAR version 2.4.2a (Dobin et al., 2013). Gene counts were derived from the number of uniquely aligned unambiguous reads by Subread:featureCount (Liao et al., 2013), version 1.4.6, with mm10 annotation gencode vM9 (Harrow et al., 2012). Batch correction was used to identify genes that were reproducibly differentially expressed genes (DEGs) across the six biological replicate comparisons of paired mutant versus control samples, by normalizing counts from featureCounts for library in R using calcNormFactors from the edgeR package (Robinson et al., 2010). Common negative binomial dispersion and empirical Bayes tagwise dispersions were estimated, and these estimations were used to a fit a negative generalized log-linear model using the edgeR package. Deviance residuals were calculated using the R package RUVseq. Upper quartile normalization was performed using betweenLaneNormalization from the EDASeq R package. Unwanted variables were then removed using RUVr from the RUVSeq R package. The number of factors of unwanted variation, k, was chosen such that k was the minimal number of factors required to produce separation in a PCA plot as described above. Batch corrected counts were used to perform DEG analysis and generate PCA plots. All gene-level transcript counts were imported into the R/Bioconductor package DESeq2 (Love et al., 2014). Transcripts with CPM >1.0 were converted into a DESeq2 dataset and then regularized log transformed using the rlog function from the DESeq2 package. Adjusted p-values for differentially expressed genes (DEGs) were determined by DESeq2 using the R stats function p.adjust using the Benjamini and Hochberg correction to determine the false discovery rate, with a >1.5-fold expression change and FDR of <0.05 required to consider a gene differentially expressed. Principle Component Analysis was performed using plotPCA from the DESeq2 package and plotted using ggplot2.

To uncover the biological significance of DEGs, network analysis was performed with the data interpretation tool Ingenuity Pathway Analysis (IPA) (Qiagen). IPA’s Ingenuity Knowledge Base uses network-eligible DEGs to generate networks and to define connections between one or more networks. Based on the number of eligible DEGs, IPA defines network scores as inversely proportional to the probability of finding the network and defines significant networks (p≤0.001). Within each network, red symbols indicate upregulated genes and green symbols indicate downregulated genes, where the color intensity represents relative degree of differential expression.

## Acknowledgements

We thank F. Mariani for critically reading the manuscript. We thank the Genome Technology Access Center in the Department of Genetics at Washington University School of Medicine for providing genomic sequencing services. Drs. David Ornitz, Cliff Tabin (obtained from Jackson laboratory), and Mark Lewandoski provided the *Dermo^Cre^*, Prx-Cre, and T-Cre mice, respectively.

## Data availability

The RNA-seq data generated during the current study are available in the Gene Expression Omnibus (GEO) repository as Series GSE143211.

## Supplemental Table Legends

**Table S1. Range of limb skeletal phenotypes in *Gmnn^f/f^*; Prx-Cre mutant animals.** (A) Frequency of each genotype in animals resulting from crosses of *Gmnn^f/f^* females and *Gmnn^+/f^*; Prx-Cre males. Mendelian ratios of animals with the *Gmnn^f/f^*; Prx-Cre (mutant) genotype were obtained. (B) Range of skeletal phenotypes in bone preparations from *Gmnn^f/f^*; Prx-Cre (mutant) and *Gmnn^+/f^*; Prx-Cre (control) animals. Age of animal at time of bone preparation and description and scoring of the number (#) of normal forelimbs, versus those with either loss or reduction of skeletal elements are indicated. (C) Summary of the frequency and range of skeletal element abnormalities observed in *Gmnn^f/f^*; Prx-Cre mutant animals.

**Table S2. Range of limb skeletal phenotypes in *Gmnn^f/f^*; *Dermo^Cre^* mutant animals.** (A) Frequency of each genotype in animals resulting from crosses of *Gmnn^f/f^* females and *Gmnn^+/f^*; *Dermo^Cre^* males. Mendelian ratios of the *Gmnn^f/f^*; *Dermo^Cre^* mutant genotype were obtained. (B) Range of skeletal phenotypes observed in bone preparations from *Gmnn^f/f^*; *Dermo^Cre^* (mutant) and *Gmnn^+/f^*; *Dermo^Cre^* (control) animals. Age of animal at time of bone preparation and description and scoring of numbers of hindlimb digits, and hindlimb skeletal structural abnormalities (reduction, loss, or curvature), and numbers of limbs exhibiting polydactyly or altered hindlimb skeletal element morphology are indicated. (C) Summary of the frequency with which polydactyly and skeletal element abnormalities were observed in *Gmnn^f/f^*; *Dermo^Cre^* mutant animals.

**Table S3. Summary of results from whole mount in situ hybridization (WISH) experiments.** (A) Pairs of somite-matched *Gmnn^f/f^*; Prx-Cre (mutant) and *Gmnn^+/f^*; Prx-Cre (control) embryos from the same timed pregnancy were analyzed in parallel by WISH for each marker for each biological replicate experiment. Summary of findings and number of biological replicate experiments conducted are shown. (B) A similar analysis was performed for *Gmnn^f/f^*; *Dermo^Cre^* (mutant) and *Gmnn^+/f^*; *Dermo^Cre^* (control) embryos. Summary of findings and the number of biological replicate experiments performed is indicated.

**Table S4. RNA-seq analysis of differentially expressed genes in *Gmnn^f/f^*; Prx-Cre mutant versus *Gmnn^+/f^*; Prx-Cre control embryonic forelimb buds.** Six pairs of somite-matched E10.5 embryos with mutant and control genotypes (with each pair obtained from the same timed pregnancy) were used to obtain forelimb buds. These were subjected to RNA-seq analysis and differentially expressed genes (DEGs) were defined as described in the methods. (A) For each DEG, the mean expression level, fold difference, p-value, and false discovery rate (FDR) are reported. (B-G) Ingenuity Pathway Analysis (IPA) of DEGs. (B-C) Class- and function-associated terms (C) and networks (D) associated with all DEGs. (D-E) Class- and function-associated terms (D) and networks (E) associated with down-regulated DEGs (lower expression in mutants versus controls). (F-G) Class- and function-associated terms (F) and networks (G) associated with up-regulated DEGs (higher expression in mutants versus controls).

## Supplemental Figure Legends

**Fig. S1. Temporal differences in *Dermo^Cre^* versus Prx-Cre expression in the fore-versus hindlimb bud.** Prx-Cre and *Dermo^Cre^* males were crossed to the R26R LacZ reporter line, and Cre expression in the developing limb buds was assessed by β-galactosidase staining of histological sections through E9.5 and E10.5 embryos.

**Fig. S2. *Gmnn^f/f^*; Prx-Cre mutant forelimbs exhibit absent and reduced skeletal elements by E14.5.** Histological comparisons of G*mnn^+/f^*; Prx-Cre (control) and *Gmnn^f/f^*; Prx-Cre (mutant) embryonic forelimbs were made by sectioning and hematoxylin and eosin staining at E14.5, 16.5, and 18.5. Mutant forelimbs exhibit loss and reduction of skeletal elements by E14.5, while differentiation of elements that remain in the mutant does not appear to be grossly impaired or delayed at E16.5-18.5.

**Fig. S3. Expression of a *β-galactosidase* reporter from the *Gmnn* locus indicates that *Gmnn* is broadly expressed in the embryo and forming limb buds.** Embryos heterozygous for the *Gmnn* knockout-first allele (*Gmnn^tm1a(KOMP)Wtsi^*), which has knock-in of a splice acceptor-*β-galactosidase* reporter cassette into the *Gmnn* gene, were used to assess *Gmnn* expression in the embryo and limb bud. Somite-matched control littermates that did not carry this allele were co-stained as controls to detect the level and pattern of *β-galactosidase* expression. Bright field images of (A) E10.5-12.5 embryos and (B) forelimb and hindlimb buds are shown, with a hindlimb section at right.

**Fig. S4. *Gmnn^f/f^*; T-Cre embryos exhibit embryonic lethality by embryonic day 9.5, with elevated apoptosis and subsequent loss of caudal tissues.** (A-B) Bright field images of (A) *Gmnn^+/f^*; T-Cre (control) and (B) *Gmnn^f/f^*; T-Cre (mutant) embryos at E9.5. As in other experimental work, somite-matched pairs of mutant and control embryos obtained from the same timed pregnancy were analyzed. (C-D) Transverse sections through E8.5 control and mutant embryos were generated at the approximate anterior (A) and posterior (P) locations shown in (C) and are indicated on the sectional view in (D) (red arrowheads). (E-O) Immunofluorescence with the antibodies indicated defined their expression in A and/or P sections of control and mutant embryos. DAPI is shown in blue and indicated antibody staining is shown in green: (E) *Brachyury* (*T/Bry*) is expressed in the trunk notochord, (F-G) *Foxa2* in the notochord, neural tube floorplate, and gut endoderm, and (H-I) *Sox1* in the neural tube. (J-K) Ki67 is a marker of all proliferating cells, (L-M) Phosphorylated histone H3 (pH3) is a marker of mitotic cells, and (N-O) cleaved, activated Caspase 3 (Caspase 3) is a marker of apoptosis. Scale bar=50µm; n=3 biological replicate experiments.

**Fig. S5. Expression of proliferative and apoptotic markers is similar in forelimb buds of *Gmnn^f/f^*; Prx-Cre mutant and *Gmnn^+/f^*; Prx-Cre control embryos.** Pairs of somite-matched mutant and control embryos from the same timed pregnancy were used to obtain E10.5 embryos and immunofluorescence on forelimb bud sections was performed with antibodies for Ki67, pH3, and cleaved Caspase 3. Scale bar=50µm.

**Fig. S6. Differences in gene expression in *Gmnn^f/f^*; Prx-Cre mutant versus *Gmnn^+/f^*; Prx-Cre control forelimb buds.** Forelimb buds were dissected from six pairs of somite-matched mutant and control E10.5 embryos (with each pair obtained from the same timed pregnancy) and subjected to RNA-seq analysis, as described in the methods. (A) Differentially expressed genes are plotted, based upon their log2 fold difference between mutant versus control expression and the FDR/adjusted p-value. (B) Genes within the ‘Gene Expression-Transcription’ term are shown. Red symbols indicate upregulated genes and green symbols indicate downregulated genes, where the color intensity represents relative degree of differential expression.

**Fig. S7. Networks associated with differentially expressed genes in *Gmnn^f/f^*; Prx-Cre mutant versus *Gmnn^+/f^*; Prx-Cre control forelimb buds.** (A-B) Selected networks identified by IPA, including DEGs that were both up- and down-regulated in mutant versus control forelimb buds. (C-D) Selected networks identified by IPA, including only DEGs that were down-regulated in mutant versus control forelimb buds. Red symbols indicate upregulated genes and green symbols indicate downregulated genes, where the color intensity represents relative degree of differential expression.

