## Supplemental Figures S1-S7 and Supplemental Tables S1-S3 for "Geminin is required for *Hox* gene regulation to pattern the developing limb"

Fig. S1

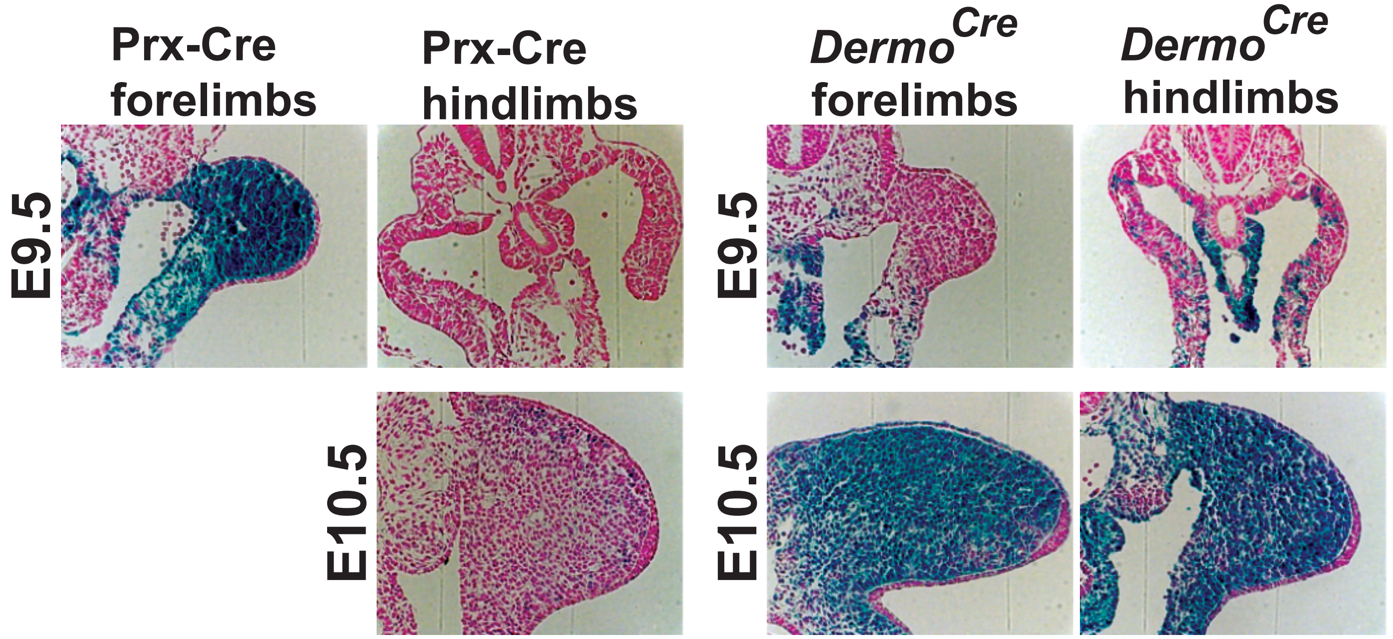
Prx-Cre forelimbs
Prx-Cre hindlimbs
*Dermo*$^{Cre}$ forelimbs
*Dermo*$^{Cre}$ hindlimbs
E9.5
E9.5
E10.5
E10.5

Fig. S2

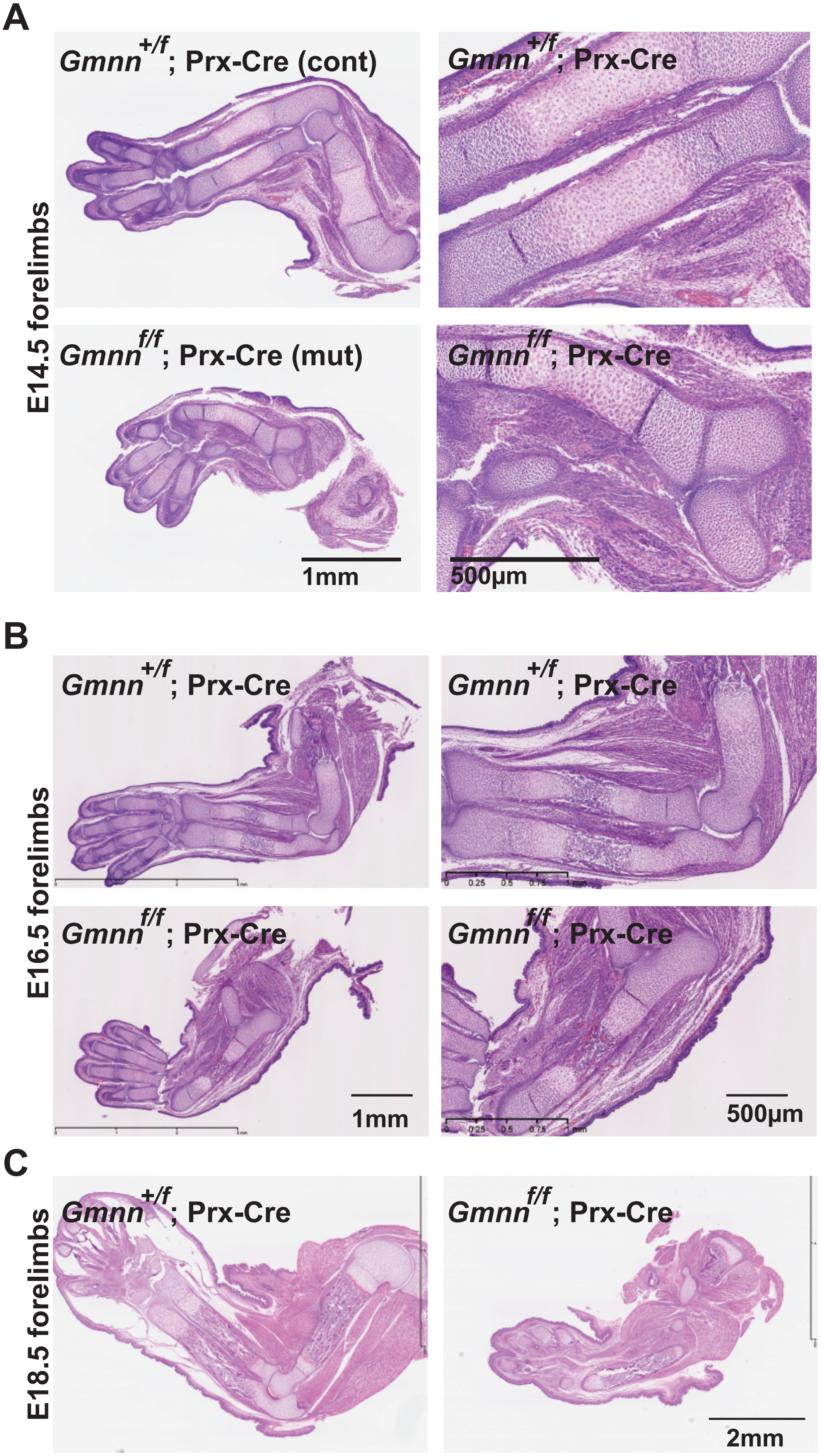
A
E14.5 forelimbs
Gmnn+/f; Prx-Cre (cont)
Gmnn+/f; Prx-Cre
Gmnnf/f; Prx-Cre (mut)
Gmnnf/f; Prx-Cre
1mm
500µm
B
E16.5 forelimbs
Gmnn+/f; Prx-Cre
Gmnn+/f; Prx-Cre
Gmnnf/f; Prx-Cre
Gmnnf/f; Prx-Cre
1mm
500µm
C
E18.5 forelimbs
Gmnn+/f; Prx-Cre
Gmnnf/f; Prx-Cre
2mm

Fig. S3

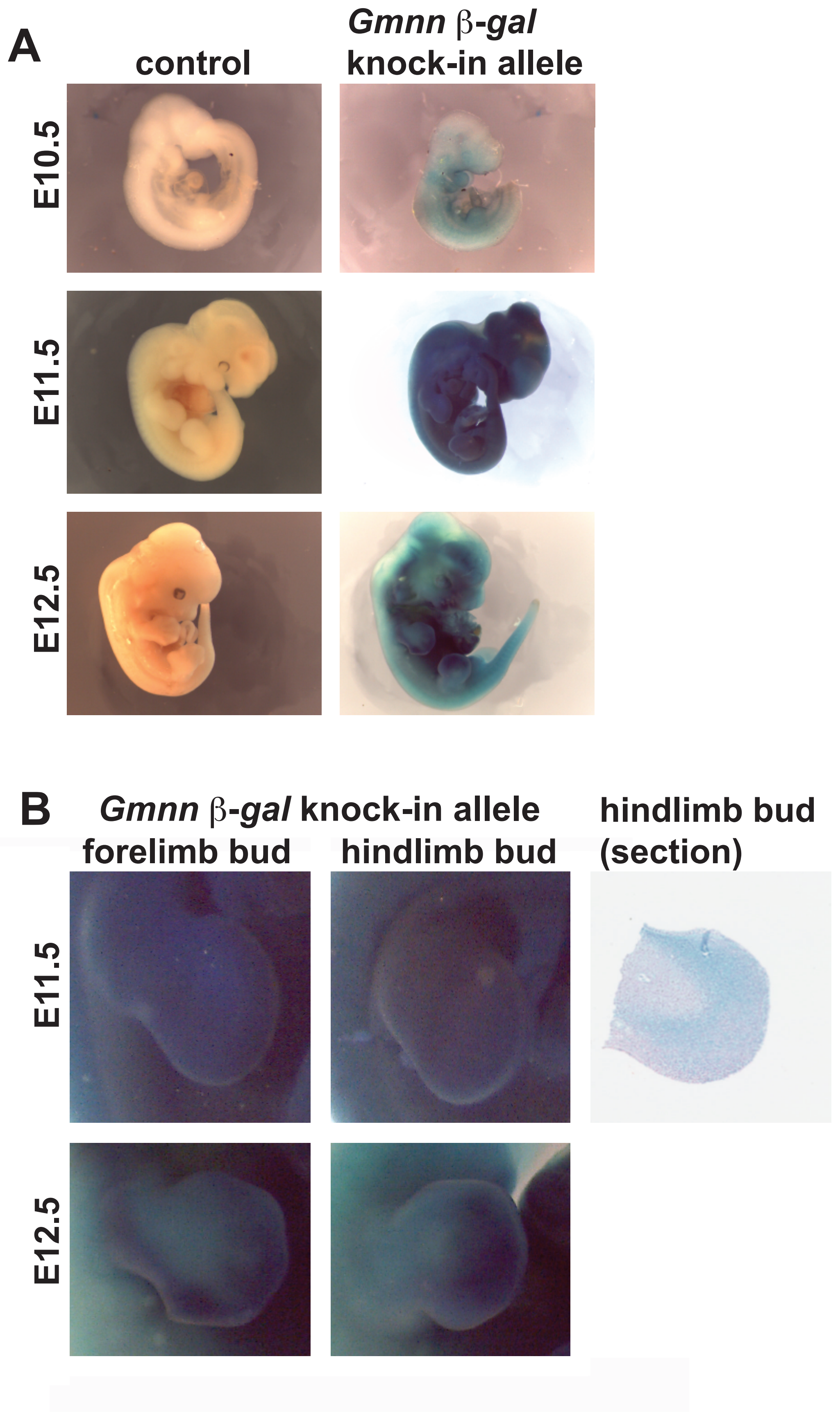
A
control
Gmnn β-gal
knock-in allele
E10.5
E11.5
E12.5
B
Gmnn β-gal knock-in allele
forelimb bud
hindlimb bud
hindlimb bud
(section)
E11.5
E12.5

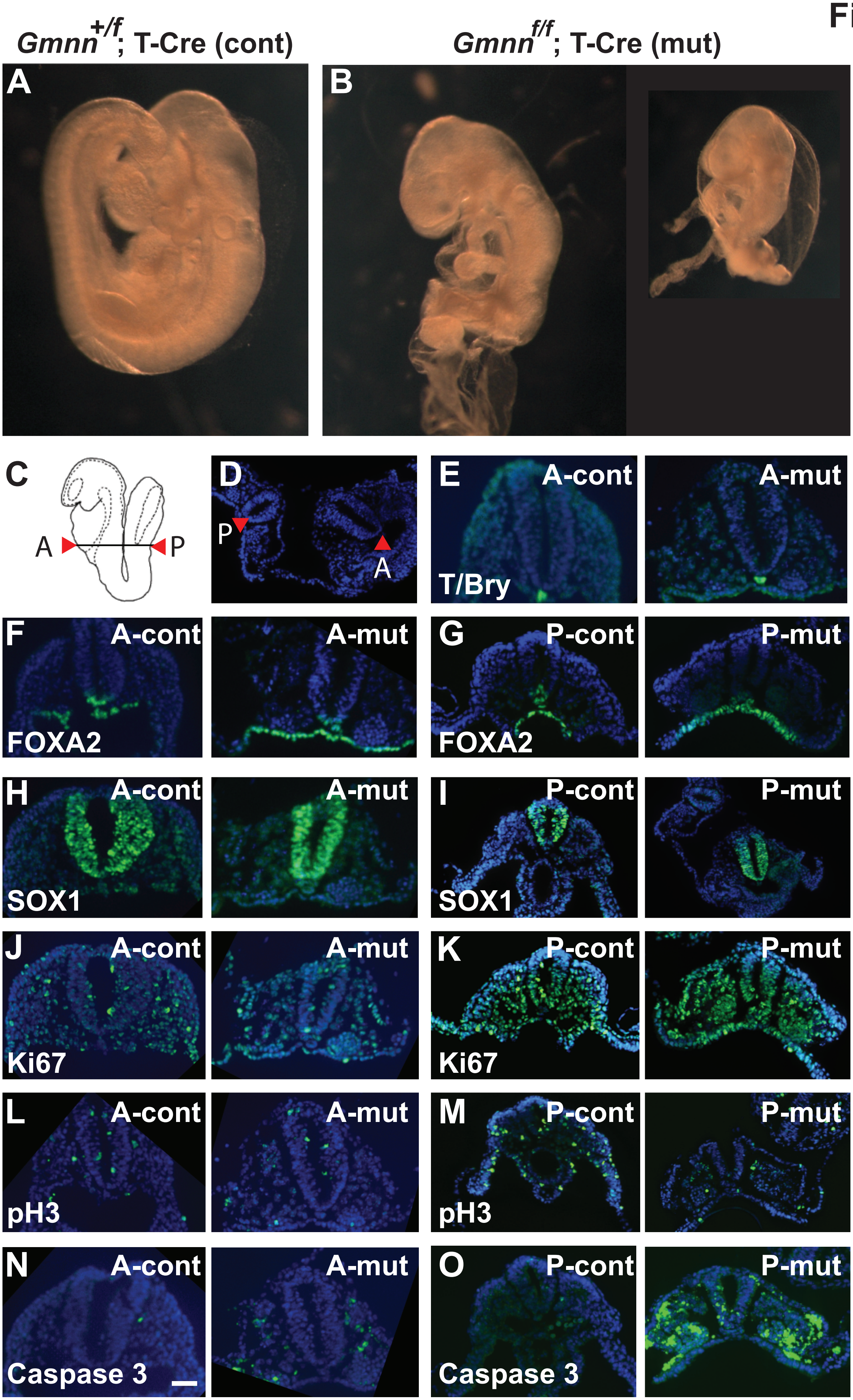

$Gmnn^{+/f}$; T-Cre (cont)
$Gmnn^{f/f}$; T-Cre (mut)
A
B
C
A
P
D
P
A
E
A-cont
A-mut
T/Bry
F
A-cont
A-mut
FOXA2
G
P-cont
P-mut
FOXA2
H
A-cont
A-mut
SOX1
I
P-cont
P-mut
SOX1
J
A-cont
A-mut
Ki67
K
P-cont
P-mut
Ki67
L
A-cont
A-mut
pH3
M
P-cont
P-mut
pH3
N
A-cont
A-mut
Caspase 3
O
P-cont
P-mut
Caspase 3

**Fig. S5**

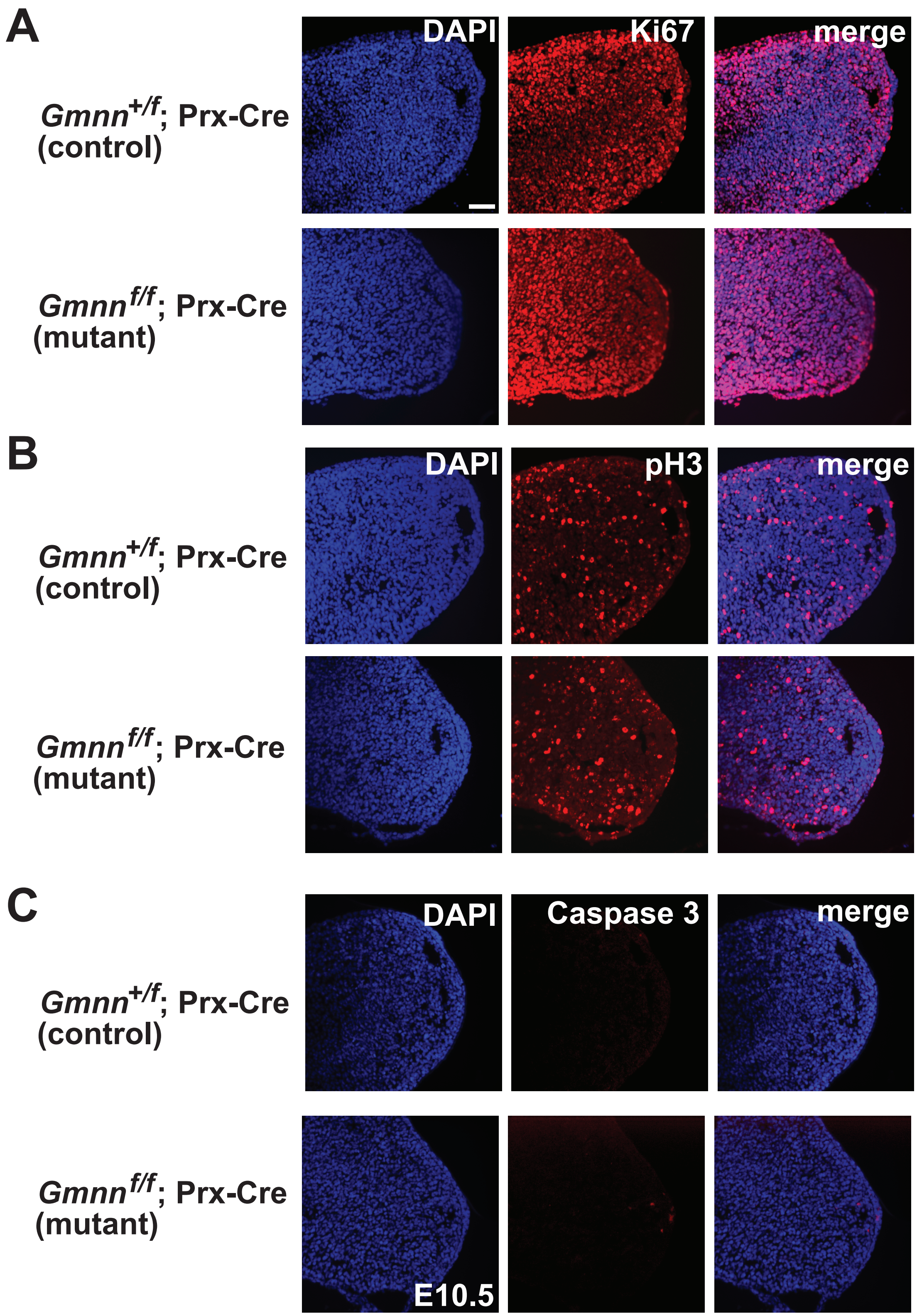

A
*Gmnn*+/f; Prx-Cre
(control)
*Gmnn*f/f; Prx-Cre
(mutant)
DAPI
Ki67
merge
B
*Gmnn*+/f; Prx-Cre
(control)
*Gmnn*f/f; Prx-Cre
(mutant)
DAPI
pH3
merge
C
*Gmnn*+/f; Prx-Cre
(control)
*Gmnn*f/f; Prx-Cre
(mutant)
DAPI
Caspase 3
merge
E10.5

**Fig. S6**

A

B

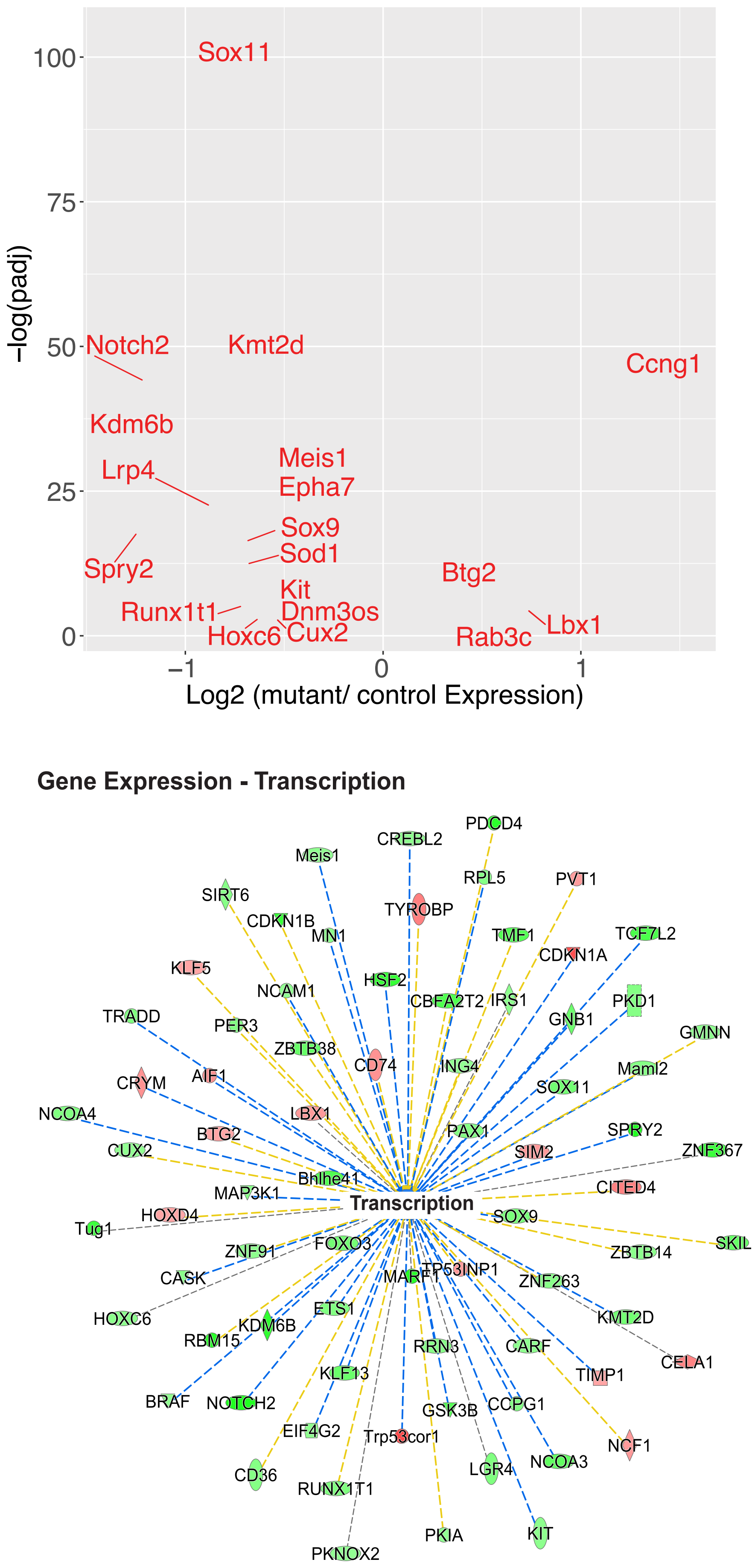

Sox11
Notch2
Kmt2d
Ccng1
Kdm6b
Lrp4
Meis1
Epha7
Sox9
Sod1
Spry2
Btg2
Kit
Runx1t1
Dnm3os
Lbx1
Hoxc6
Cux2
Rab3c
100
75
50
25
0
−1
0
1
−log(padj)
Log2 (mutant/ control Expression)
Gene Expression - Transcription
Transcription
PDCD4
CREBL2
Meis1
RPL5
PVT1
SIRT6
TYROBP
CDKN1B
MN1
TMF1
TCF7L2
CDKN1A
KLF5
HSF2
PKD1
NCAM1
CBFA2T2
IRS1
TRADD
GNB1
GMNN
PER3
ZBTB38
CD74
ING4
Maml2
CRYM
AIF1
SOX11
NCOA4
LBX1
PAX1
SPRY2
BTG2
SIM2
ZNF367
CUX2
Bhlhe41
CITED4
MAP3K1
HOXD4
SOX9
Tug1
SKIL
ZNF91
FOXO3
ZBTB14
CASK
TP53INP1
MARF1
ZNF263
HOXC6
ETS1
KMT2D
KDM6B
RBM15
RRN3
CARF
CELA1
KLF13
TIMP1
BRAF
NOTCH2
GSK3B
CCPG1
EIF4G2
Trp53cor1
NCF1
LGR4
NCOA3
CD36
RUNX1T1
PKIA
KIT
PKNOX2

### Fig. S7

**A** **All DEGs: Cellular Development, Organismal Development, Reproductive System Development and Function**

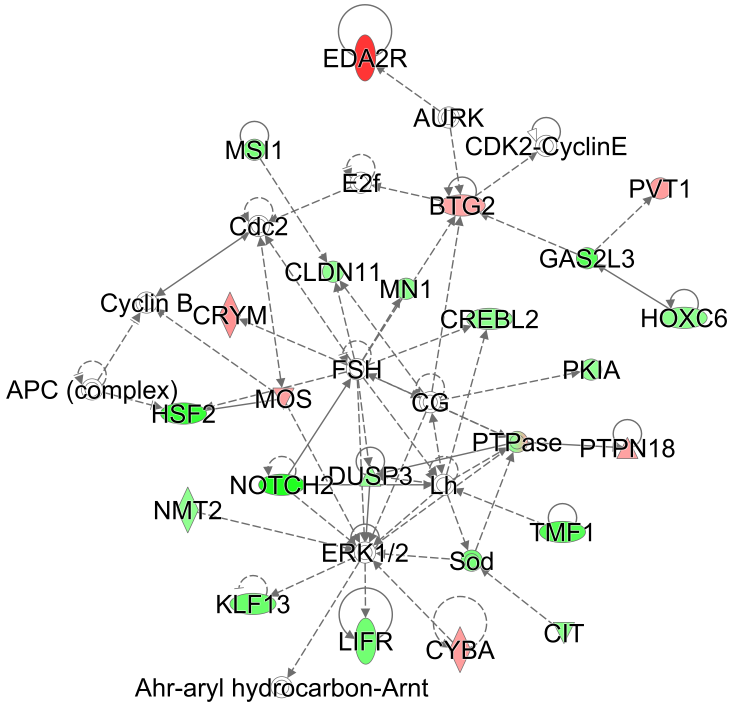


**B** **All DEGs: Cell Death and Survival, Cellular Development, Cellular Growth and Proliferation**

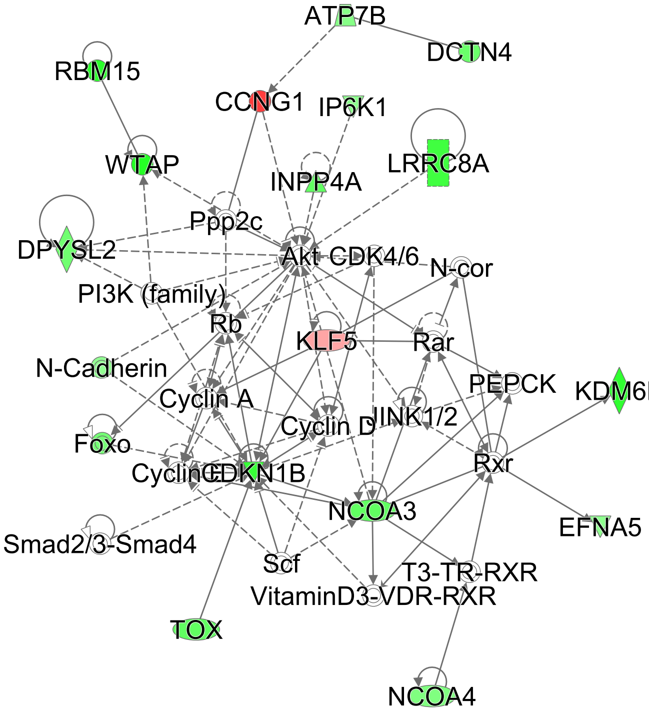


**C** **Downregulated DEGs: Connective Tissue Disorders, Developmental Disorder, Organismal Injury and Abnormalities**

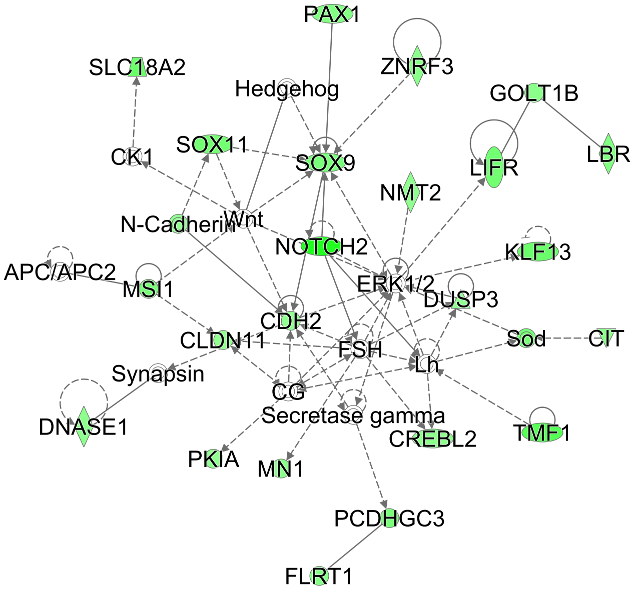


**D** **Downregulated DEGs: Cellular Assembly and Organization, Cellular Function and Maintenance, Embryonic Development**

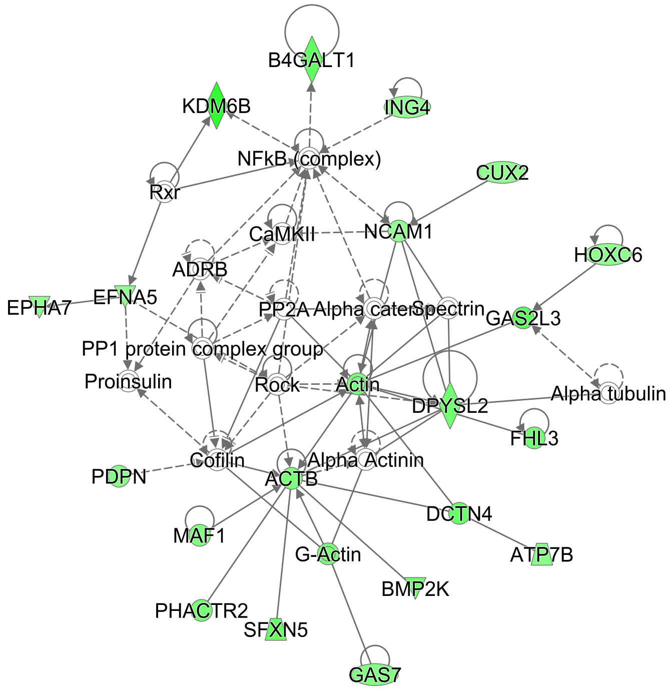

**Table S1. Range of Phenotypes in *Gmnn*$^{f/f}$; Prx-Cre animals**

**A. Frequency of each genotype in animals resulting from matings of male *Gmnn*$^{+/f}$; Prx-Cre to female *Gmnn*$^{f/f}$**

| Genotype | n | % |
|---|---|---|
| *Gmnn*$^{f/f}$; Prx-Cre (mut) | 27 | 25.23 |
| *Gmnn*$^{+/f}$; Prx-Cre (cont) | 35 | 32.71 |
| *Gmnn*$^{+/f}$ | 21 | 19.63 |
| *Gmnn*$^{f/f}$ | 24 | 22.43 |
| **n=** | **107** | |

**B. Range of skeletal phenotypes observed in bone preparations from mutant and control animals**

| ***Gmnn*$^{f/f}$; Prx-Cre (mut) genotype (n=10)** | | | | | |
|---|---|---|---|---|---|
| **Age at time of bone preparation** | **Forelimb skeletal elements** | **# normal forelimb** | **# forelimb with all elements present but shortened** | **# forelimb missing humerus** | **# forelimb missing radius and/or ulna** |
| E18.5 | One side very short shoulder girdle, missing humerus, missing either radius or ulna; other side very short shoulder girdle, missing humerus, missing either radius or ulna. | 0 | 0 | 2 | 2 |
| P5 | One side short shoulder girdle, no humerus, missing either radius or ulna; other side very short shoulder girdle missing humerus, missing either radius or ulna. | 0 | 0 | 2 | 2 |
| P25 | Both sides, all elements present but shorter. | 0 | 2 | 0 | 0 |
| P25 | One side short shoulder girdle, curved humerus; other side very short radius or ulna, but both present. | 0 | 2 | 0 | 0 |
| P25 | One side has all elements present but shorter. Other side missing humerus, with curved either radius or ulna and one element missing. | 0 | 1 | 1 | 1 |
| P25 | Overall much shorter but all elements are present on both sides, one side smaller shoulder girdle; the other side all elements are just shorter. | 0 | 2 | 0 | 0 |
| P25 | One side has all elements present but shorter. One side missing humerus, has curved either radius or ulna and one missing. | 0 | 1 | 1 | 1 |
| P25 | One side severe very short shoulder girdle, missing humerus, missing either radius or ulna. Other side all elements are present but shorter. | 0 | 1 | 1 | 1 |
| P25 | One side very short shoulder girdle , missing humerus, missing either radius/ulna. Other side short but all elements present. | 0 | 1 | 1 | 1 |
| P25 | One side very short shoulder girdle, missing humerus, missing either radius or ulna. Other side all elements present but shortened. | 0 | 1 | 1 | 1 |

| ***Gmnn*$^{+/f}$; Prx-Cre (control) genotype (n=6)** | | | | | |
|---|---|---|---|---|---|
| **Age at time of bone preparation** | **Forelimb skeletal elements** | **# normal forelimb** | **# forelimb with all elements present but shortened** | **# forelimb missing humerus** | **# forelimb missing radius and/or ulna** |
| E18.5 | All elements present, normal morphology. | 2 | 0 | 0 | 0 |
| P5 | All elements present, normal morphology. | 2 | 0 | 0 | 0 |
| P25 | All elements present, normal morphology. | 2 | 0 | 0 | 0 |
| P25 | All elements present, normal morphology. | 2 | 0 | 0 | 0 |
| P25 | All elements present, normal morphology. | 2 | 0 | 0 | 0 |
| P25 | All elements present, normal morphology. | 2 | 0 | 0 | 0 |

**C. Summary of range of skeletal element abnormalities observed in *Gmnn*$^{f/f}$; Prx-Cre mutant animals**

*Gmnn*$^{+/f}$; Prx-Cre control n=6; 100% normal skeletal morphology
*Gmnn*$^{f/f}$; Prx-Cre mutants n=10; 0% normal skeletal morphology

**In the mutant animals:**

30% (3) had shortening of some or all of the forelimb skeletal elements, but all elements present.
50% (5) had one limb with shortened elements all present, while the humerus and radius or ulna were missing on the other limb.
20% (2) had missing humerus and either radius or ulna in both forelimbs.

Table S2. Range of Phenotypes in *Gmnn*$^{f/f}$ *; Dermo*$^{Cre}$ animals

A. Frequency of each genotype in animals resulting from matings of male *Gmnn*$^{+/f}$ *; Dermo*$^{Cre}$ to female *Gmnn*$^{f/f}$

| Genotype | n | % |
|---|---|---|
| *Gmnn*$^{f/f}$*; Dermo*$^{Cre}$ | 32 | 24.62 |
| *Gmnn*$^{+/f}$*; Dermo*$^{Cre}$ | 42 | 32.31 |
| *Gmnn*$^{+/f}$ | 24 | 18.46 |
| *Gmnn*$^{f/f}$ | 32 | 24.62 |
| n= | 130 | |

B. Range of skeletal phenotypes observed in bone preparations from mutant and control animals

| *Gmnn*$^{f/f}$ *; Dermo*$^{Cre}$ (mut) genotype (n=26) | | | | | | |
|---|---|---|---|---|---|---|
| Age at time of bone preparation | Left foot (# digits) | Right foot (# digits) | Left hindlimb skeletal element alterations | Right hindlimb skeletal element alteration | # hindlimbs with polydactyly | # hindlimbs with altered skeletal element morphology |
| E18.5 | 6 | 6 | normal | normal | 2 | 0 |
| E18.5 | 6 | 6 | normal | normal | 2 | 0 |
| E18.5 | 6 | 6 | normal | normal | 2 | 0 |
| E18.5 | 6 | 6 | normal | normal | 2 | 0 |
| E18.5 | 6 | 6 | normal | normal | 2 | 0 |
| E18.5 | 6 | 6 | normal | normal | 2 | 0 |
| E18.5 | 6 | 6 | normal | normal | 2 | 0 |
| E18.5 | 6 | 6 | normal | normal | 2 | 0 |
| E18.5 | 6 | 6 | normal | normal | 2 | 0 |
| P5 | 7 | 6 | normal | normal | 2 | 0 |
| adult | 6 | 6 | normal | normal | 2 | 0 |
| adult | 6 | 6 | shortened and curved | shortened and curved | 2 | 2 |
| adult | 5 | 5 | very short femur, missing tibia/fibula | shortened and curved | 0 | 2 |
| adult | 6 | 7 | normal | normal | 2 | 0 |
| adult | 7 | 6, syndactyly | normal | normal | 2 | 0 |
| adult | 6 | 6 | shortened and curved | curved | 2 | 2 |
| adult | 6 | 6 | shortened | shortened | 2 | 2 |
| adult | 6 | 7, syndactyly | curved | curved | 2 | 2 |
| adult | 7, syndactyly | 7, syndactyly | shortened and curved | normal | 2 | 1 |
| adult | 7 | 6, syndactyly | normal | shortened | 2 | 1 |
| adult | 6 | 7, syndactyly | normal | shortened and curved | 2 | 1 |
| adult | 6, syndactyly | 6, syndactyly | normal | shortened and curved | 2 | 1 |
| adult | 6 | 6 | shortened and curved | shortened and curved | 2 | 2 |
| adult | 7 | 6 | shortened and curved | shortened and curved | 2 | 2 |
| adult | 5 | 6, syndactyly | normal | normal | 1 | 0 |
| adult | 5 | 5 | very short | normal | 0 | 1 |

| *Gmnn*$^{+/f}$*; Dermo*$^{Cre}$ (control) genotype (n=9) | | | | | | |
|---|---|---|---|---|---|---|
| Age at time of bone preparation | Left foot (# digits) | Right foot (# digits) | Left hindlimb skeletal element alterations | Right hindlimb skeletal element alteration | # hindlimbs with polydactyly | # hindlimbs skeletal element morphology |
| E18.5 | 5 | 5 | normal | normal | 0 | 0 |
| E18.5 | 5 | 5 | normal | normal | 0 | 0 |
| P5 | 5 | 5 | normal | normal | 0 | 0 |
| P5 | 5 | 5 | normal | normal | 0 | 0 |
| P5 | 5 | 5 | normal | normal | 0 | 0 |
| P5 | 5 | 5 | normal | normal | 0 | 0 |
| adult | 5 | 5 | normal | normal | 0 | 0 |
| adult | 5 | 5 | normal | normal | 0 | 0 |
| adult | 5 | 5 | normal | normal | 0 | 0 |

**C. Summary of range of hindlimb abnormalities observed in *Gmnn*$^{f/f;}$ *Dermo*$^{Cre}$ mutant animals**

**Polydactyly**

92% (24/26; n=26) of *Gmnn*$^{f/f}$*; Dermo*$^{Cre}$ (mut) have polydactyly, almost always in both hind feet.
9/52 (17%) of hind feet also exhibited syndactyly uni- or bilaterally
Polydactyly included hind limbs with either 6 (38/52=73%) or 7 (9/52=17.3%) digits.
Forelimbs never exhibited polydactyly.
In several cases, polydactyly was not present, but skeletal elements were morphologically abnormal.
*Gmnn*$^{+/f}$*; Dermo*$^{Cre}$ control n=9 (0% abnormal).

**Skeletal element alterations**

Skeletal element alterations were not evident at E18.5 or P5 but were apparent in some adults.
*Gmnn*$^{f/f}$*; Dermo*$^{Cre}$ (mut), n=16 scored as adults.
Of these 32 hindlimbs, 59% have one or more skeletal elements that are either curved, shortened, or both.
87.5% have hindlimb skeletal element abnormalities visible as adults.
*Gmnn*$^{+/f}$*; Dermo*$^{Cre}$ (control) n=9 (0% abnormal).

**Table S3. Summary of whole mount in situ analysis of Gmnnf/f; Prx-Cre and Gmnnf/f; DermoCre mutant embryos**

| **A.** | **Gene expression in *Gmnn*$^{f/f}$ ; Prx-Cre mutant versus *Gmnn*$^{+/f}$ ; Prx-Cre control forelimb buds (each analysis was performed by in parallel WISH analysis of pairs of somite-matched mutant and control embryos with these genotypes that were obtained from the same timed pregnancy).** | **# pairs of mutant and control embryos** |
|---|---|---|
| *Hoxd10* | Expression domain in mutant forelimb buds is expanded proximally, is more symmetric along the anteroposterior axis, and has less distinct borders than the control. | 3 |
| *Hoxd11* | Expression domain in mutant forelimb buds is expanded proximally and is more symmetric along the anteroposterior axis than the control. | 6 |
| *Hoxd12* | Expression domain in mutant forelimb buds is expanded proximally and is more symmetric along the anteroposterior axis than the control. | 3 |
| *Hoxd13* | Expression domain in mutant forelimb buds is expanded proximally versus control. | 4 |
| *Hoxa13* | Expression domain in mutant forelimb buds is expanded proximally versus control. | 4 |
| *Ptch1* | Expression domain in mutant forelimb buds is expanded anteriorly versus control. | 4 |
| *Hand2* | Expression domain similar in mutant forelimb buds and control. | 3 |
| *Shh* | Expression domain similar in mutant forelimb buds and control. | 3 |
| *Fgf8* | Expression domain similar in mutant forelimb buds and control. | 3 |

| **B.** | **Gene expression in *Gmnn*$^{f/f}$ *; Dermo*$^{Cre}$ mutant versus *Gmnn*$^{+/f}$ *; Dermo*$^{Cre}$ control hindlimb buds (each analysis was performed by in parallel WISH analysis of somite-matched pairs of mutant and control embryos with these genotypes that were obtained from the same timed pregnancy).** | **# pairs of mutant and control embryos** |
|---|---|---|
| *Hoxd10* | Expression domain similar in mutant and control. | 2 |
| *Hoxd11* | Expression domain similar in mutant and control. | 5 |
| *Hoxd12* | Expression domain similar in mutant and control at E11.5; mutant has expanded expression to the anterior boundary of the hindlimb bud at E12.5. | 3 |
| *Hoxa13* | Expression domain similar in mutant and control. | 4 |
| *Hoxd13* | Mutant has ectopic domain of expression in the anterior hindlimb bud at low penetrance (1/8 embryos assayed). | 8 |
| *Ptch1* | Mutant has ectopic domain of expression in the anterior hindlimb bud with partial penetrance (3/4 embryos assayed). | 4 |
| *Shh* | Mutant has ectopic domain of expression in the anterior hindlimb bud with partial penetrance (3/8 embryos assayed). | 8 |
| *Hand2* | Expression domain similar in mutant and control. | 4 |
| *Fgf8* | Expression domain similar in mutant and control. | 3 |
